# Resolving cDC2 heterogeneity across human cancer atlases

**DOI:** 10.1101/2025.09.08.674999

**Authors:** Nikita Rosendahl, Xiaohan Xu, Norman Teik-Wei Yap, Zewen Kelvin Tuong, Kristen J Radford

## Abstract

Dendritic cells (DCs) are vital in driving effective anti-tumour immune responses. Multiple DC subpopulations have been identified in human tumours, however their alignment, phenotype and function across tumour types have not been comprehensively addressed. Here, we curated single-cell RNA sequencing datasets to generate an extensive integrated atlas of myeloid antigen presenting cells (APCs) across different human cancer types. We confirm that cDC1, cDC2 and mregDC represent the major DC populations present in human tumours, and that these populations are highly conserved regardless of cancer type. A distinct DC3 population was not identified within the atlas, suggesting these cells are less distinguishable in tumour tissue. The cDC2 compartment resolved into two subpopulations, a minor population that aligned tumour associated CD207^+^CD1A^+^ cDC2 with splenic cDC2A, and a predominate subset aligning with cDC2B. This integrated atlas serves as a powerful resource enabling deeper investigation into the functions of tumour-associated DC diversity, phenotype and function, across a range of human cancer types.

## Background

Dendritic cells (DCs) are critical in generating effective anti-tumour immunity, while also driving immune tolerance to maintain homeostasis. Within the tumour microenvironment (TME), DCs ingest and process tumour-associated antigens before migrating to neighbouring lymph nodes, where they present these antigens to T cells to initiate and drive immune responses.^1^ DC also play a critical role in the activation and maintenance of T cells within the TME. Single-cell transcriptomics of human tissues, including many types of cancer, has revealed significant heterogeneity within the DC compartment. DC diversity and their contextual environment are critical determinants of immune activation or regulation, and an in-depth understanding is essential for enhancing anti-tumour immunity. However, the rarity of DC, differences in clustering, and variations in annotations have made comparisons across datasets and tumour types challenging.

Conventional DCs (cDCs) arise from a common DC progenitor (CDP) through precursor cDCs (pre-cDCs) in an FMS-like tyrosine kinase 3 (FLT3)-dependent manner. Pre-cDCs are pre-committed as either pre-cDC1s or pre-cDC2s, which develop into the cDC1 and cDC2 subsets.^2^ The cDC1 subset is highly conserved across tissues and species and is critical for tumour immunity and for the efficacy of most immunotherapies in murine models. In human cancers, cDC1s are rare, but their presence is associated with good prognosis and responses to immunotherapy. By contrast, the cDC2 subset, defined by expression of CD1c, is more abundant in tumour tissue but represents a heterogeneous population, with as many as six different subpopulations described.^1,3,4^

In human blood, the cDC2 population has been segregated into CD5^+^ cDC2 and CD14^+^ CD163^+^ DC3, with the latter being defined by an inflammatory signature that overlaps with monocytes.^2,5^ DC3 are proposed to develop from a distinct bone marrow precursor to monocytes and cDC2,^6–8^ however their precise origins remain unclear. DC3-like cells (CD14^+^ DC) have been reported previously in several human cancer types including ovarian cancer ascites,^9^ melanoma,^10^ and breast cancer.^6^ Adding further complexity, monocyte-derived DC (MoDC) can also directly develop from monocytes under inflammatory conditions and these also display overlapping signatures, phenotype, and functional features with monocytes, cDC2, and DC3.^11^

In human tumours cDC2 have been segregated into numerous subpopulations and states,^1,3,4^ including ISG^+^ DCs, a population enriched in interferon-stimulated genes (ISG^+^ DC), and a population enriched in Langerhans cell-associated markers *CD1A* and *CD207* (CD207^+^ DC). How these different cDC2 subtypes relate to each other and align across different datasets, whether they represent unique ontogenetic lineages or maturation states, and whether they have specialised roles in tumour immunity remains to be established.

Another distinct cluster of DC, characterised by a unique signature, termed ‘mature DCs enriched in immunoregulatory molecules’ (mregDC, also known as migratory DC, mature DC, CCR7^+^ DC, and LAMP3^+^ DC) is also present in human tumours.^2^ This DC subtype appears to be conserved across human cancer types and can also be found in other tissues in both inflammatory and homeostatic contexts.^1,2^ The mregDC population is characterised by expression of genes involved in both DC maturation and immunoregulation, and can be derived from both cDC1 and cDC2 following the uptake of dead cells.^1^ As such, mregDCs are considered to represent a maturation state rather than an ontogenetically distinct DC lineage. Whether putative immunogenic versus regulatory functions differ across cancer types, or can be uncoupled to enhance tumour immunity, is yet to be resolved.

Consolidating DC subtypes and inferring their roles across cancers using transcriptomic datasets has been constrained due to the rarity of DCs within tumours and the limited sample sizes of available datasets. Drawing conclusions from small populations can be problematic, as low cell counts and limited sample sizes increase the risk of sampling bias, loss of rare subpopulations, or artificial cluster distinctions driven by technical noise rather than true biological differences.

Here, to more clearly address heterogeneity in myeloid antigen-presenting cells (APCs) in cancer tissue, we curated an extensive scRNA-seq integrated atlas of myeloid APCs consisting of 498,023 cells from 589 samples across 36 publicly available datasets. We focus on the annotation of DCs, but broadly included other *HLA-DR*-expressing myeloid cells, including monocytes and macrophages. The curated atlas includes ∼30,000 DCs, covering 12 cancer types (including both primary and metastatic samples) along with healthy tissue from six sites. This atlas provides a unique resource for in-depth characterisation of DC phenotype, maturation, and the heterogeneity between cancer types compared with healthy tissue. Using this resource, we have confirmed that cDC1, cDC2 and mregDC represent the major DC subtypes consistently present across human tumour tissues, and that their phenotype remains relatively conserved regardless of tissue or cancer type. The DC3 signature did not identify a unique population within tumour tissue but overlapped with monocytes and cDC2. Within the cDC2 compartment, clear populations aligning to cDC2A and cDC2B subsets were identified, with the cDC2A population also aligning with the CD207^+^ DC state. This atlas can be used in future for deeper interrogation of DC diversity, phenotype, and function across different cancer types.

## Results

### Curation of a single-cell atlas of myeloid APCs in human cancer

We curated an extensive single-cell atlas of myeloid APCs in human tumours using publicly available datasets, integrating datasets indexed on the DISCO database^12^ as well as highly-cited and recent studies not yet indexed on DISCO (**Table 1**). After extensive quality control checks and pre-preprocessing (see methods), our curated atlas covers 12 major cancer types (**Figure 1a**, **Table 1**), including breast cancer (BC), colorectal carcinoma (CRC), gastric adenocarcinoma (GAC), glioblastoma (GBM), hepatocellular carcinoma (HCC), high-grade serous ovarian carcinoma (HGSOC), head and neck squamous cell carcinoma (HNSCC), intrahepatic cholangiocarcinoma (iCCA), melanoma (MEL), nasopharyngeal carcinoma (NPC), non-small cell lung cancer (NSCLC), and pancreatic ductal adenocarcinoma (PDAC). Sample types included primary tumour (cancerous tissue taken from the primary site, including recurrence at the primary site), metastatic tumour (cancerous tissue taken from sites of distant metastases), and in the case of HGSOC, malignant ascites. We also retained single-cell data from healthy tissue, including from breast, colon, liver, lung, lymph nodes, and ovary. Most samples came from treatment naïve patients; however, some samples were post-treatment (information contained in atlas metadata, summarised in **Table 1**).

**Figure 1:**
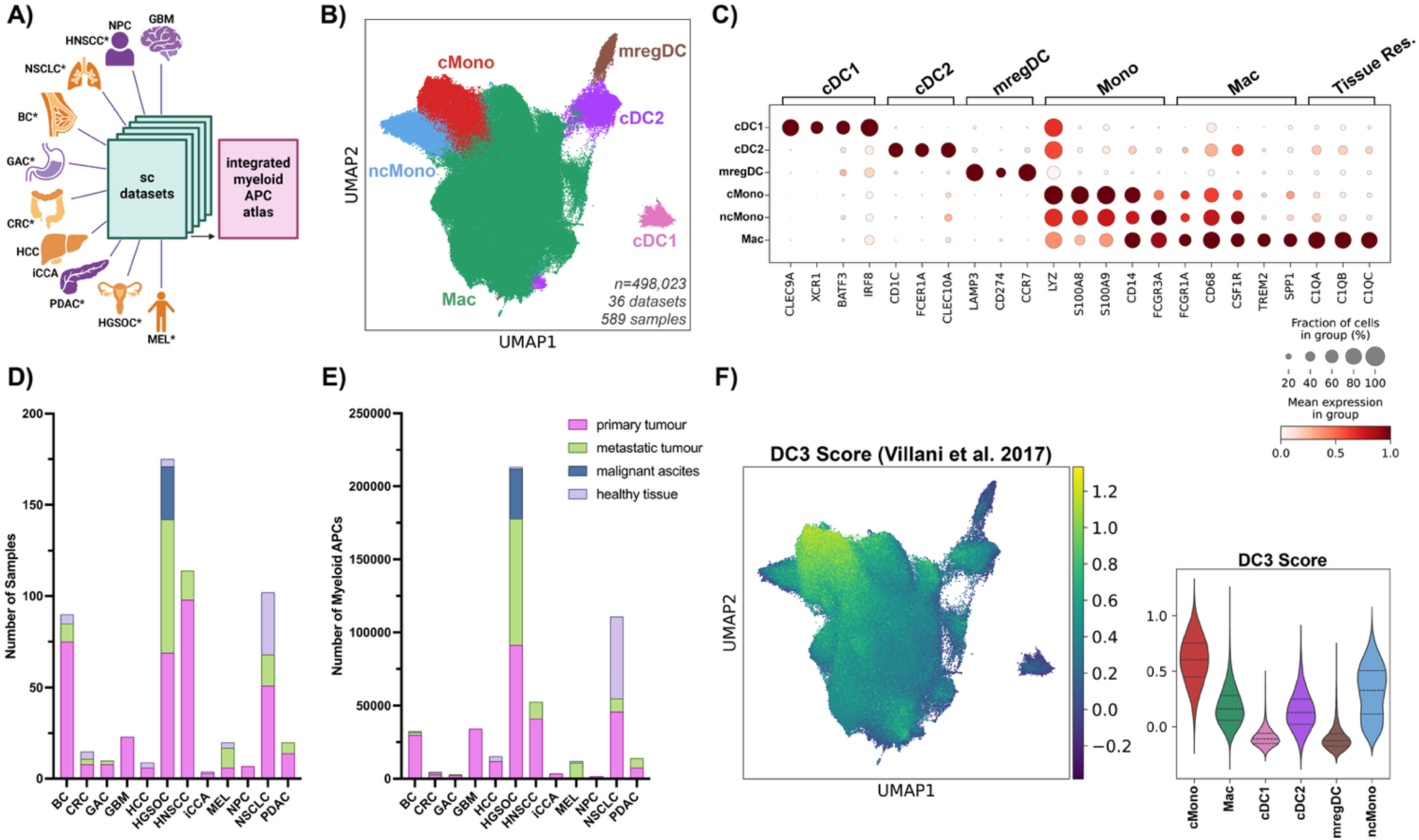
Myeloid APC single-cell atlas in human cancer. **(A)** Myeloid APCs from 37 publicly available single-cell RNA sequencing datasets, covering 12 cancer types, were integrated into a single atlas. Icons/labels indicate included primary tumour types. Orange indicates organs with additional healthy samples, *indicates cancer types with metastatic samples (various sites). Note, healthy controls for melanoma cases are uninvolved lymph nodes. Schematic created with BioRender.com. **(B)** UMAP of myeloid APC atlas, showing annotations by colour. **(C)** Select canonical markers for each cluster. Dot size indicates percent of expressing cells, colour represents expression level. **(D-E)** The final atlas contained 498,023 cells from 589 samples, and included data from 12 cancer types, with a mix of primary and metastatic tumours, and malignant ascites. Atlas also includes various healthy tissues. Plots summarise the number of cells **(D)** and number of samples **(E)** by cancer and sample type. **(F)** DC3 signature published by Villani et al^5^ was overlaid over the atlas and mean enrichment score per cell presented as a violin plot grouped by subset, lines show quartiles. Detailed statistics in Supplementary Table 3. Abbreviations: high-grade serous ovarian carcinoma (HGSOC), melanoma (MEL), breast cancer (BC), non-small cell lung cancer (NSCLC), colorectal carcinoma (CRC), hepatocellular carcinoma (HCC), intrahepatic cholangiocarcinoma (iCCA), nasopharyngeal carcinoma (NPC), head and neck squamous cell carcinoma (HNSCC), pancreatic ductal adenocarcinoma (PDAC), gastric adenocarcinoma (GAC), glioblastoma (GBM), cMono: classical monocytes, ncMono: non-classical monocytes, Mac: Tissue-resident macrophage.

**Table 1.** Summary of datasets included in the single-cell atlas of myeloid APCs in human cancer. In most cases ‘Dataset ID’ refers to a GSE accession number from the Gene expression omnibus,^42^ alternatively two samples have a ‘Dataset ID’ starting with ‘PRJ’ referring to the Chinese National Genomics Data Center, and one sample has a PubMed ID (starting with ‘PMID’). Where available, the published study associated with each dataset is provided along with the cancer type of the samples. For sample type, shading indicates whether the corresponding sample type was included in the atlas, number indicates the number of samples included. Patient treatment is recorded based on details provided in the associated studies or on the repository, and in most cases refers only to ongoing or recent treatment at the time of sample collection. IO: immunotherapy, CTX: chemotherapy, TT: Targeted therapy. ‘/’ indicates a mix of treatments among samples or combination treatments. Not specified listed for datasets where no treatment information provided.

| Accession code/ Dataset ID | Associated Study | Cancer Type | Sample Type |  |  |  | Patient Treatment |
| --- | --- | --- | --- | --- | --- | --- | --- |
|  |  |  | Primary tumour | Metastatic tumour | Healthy tissue | Malignant ascites |  |
| GSE184880 | Xu et al<sup>43</sup> | High-grade serous ovarian carcinoma (HGSOC) | 7 |  | 4 |  | Treatment naïve |
| GSE213243 | Ren et al<sup>44</sup> | High-grade serous ovarian carcinoma (HGSOC) | 1 |  |  | 1 | CTX |
| GSE217517 | Hippen et al<sup>45</sup> | High-grade serous ovarian carcinoma (HGSOC) | 7 |  |  |  | Treatment naïve |
| GSE180661 | Vazquez-Garcia et al<sup>17</sup> | High-grade serous ovarian carcinoma (HGSOC) | 45 | 69 |  | 20 | Treatment naïve |
| PRJCA005422 | Zheng et al<sup>18</sup> | High-grade serous ovarian carcinoma (HGSOC) | 9 | 4 |  | 8 | Treatment naïve |
| GSE200218 | Biermann et al<sup>46</sup> | Melanoma (MEL) |  | 5 |  |  | Treatment naïve |
| GSE215120 | Zhang et al<sup>47</sup> | Melanoma (MEL) | 6 | 1 |  |  | Treatment naïve |
| PRJNA907381 | He et al<sup>48</sup> | Melanoma (MEL) |  | 5 | 3 |  | Treatment naïve/ IO |
| GSE161529 | Pal et al<sup>49</sup> | Breast Cancer (BC) | 27 | 3 | 4 |  | Treatment naïve |
| GSE176078 | Wu et al<sup>50</sup> | Breast Cancer (BC) | 22 |  |  |  | Treatment naïve/ CTX |
| GSE195861 | Tokura et al<sup>51</sup> | Breast Cancer (BC) | 9 | 5 | 1 |  | Treatment naïve |
| GSE199515 | N/A | Breast Cancer (BC) | 2 |  |  |  | not specified |
| GSE225600 | Liu et al<sup>52</sup> | Breast Cancer (BC) | 3 | 2 |  |  | Treatment naïve |
| GSE162498 | Gueguen et al<sup>53</sup> | Non-small cell lung cancer (NSCLC) | 11 |  | 2 |  | Treatment naïve |
| GSE131907 | Kim et al<sup>54</sup> | Non-small cell lung cancer (NSCLC) | 15 | 17 | 11 |  | Treatment naïve |
| GSE154826 | Leader et al<sup>55</sup> | Non-small cell lung cancer (NSCLC) | 25 |  | 21 |  | Treatment naïve |
| PMID32561858 | Qian et al<sup>16</sup> | Breast Cancer (BC) | 12 |  |  |  | Treatment naïve |
|  |  | Colorectal Carcinoma (CRC) | 7 |  | 4 |  | Treatment naïve |
| GSE112271 | Losic et al<sup>56</sup> | Hepatocellular carcinoma (HCC) | 2 |  |  |  | Treatment naïve |
| GSE189903 | Ma et al<sup>57</sup> | Hepatocellular carcinoma (HCC) | 4 |  | 3 |  | not specified |
|  |  | Intrahepatic cholangiocarcinoma (iCCA) | 3 |  | 1 |  | not specified |
| GSE162025 | Liu et al<sup>58</sup> | Nasopharyngeal carcinoma (NPC) | 7 |  |  |  | Treatment naïve |
| GSE139324 | Cillo et al<sup>59</sup> | Head and neck squamous cell carcinoma (HNSCC) | 22 |  |  |  | Treatment naïve |
| GSE164690 | Kurten et al<sup>60</sup> | Head and neck squamous cell carcinoma (HNSCC) | 18 |  |  |  | Treatment naïve |
| GSE173468 | N/A | Head and neck squamous cell carcinoma (HNSCC) | 14 |  |  |  | not specified |
| GSE188737 | Quah et al<sup>61</sup> | Head and neck squamous cell carcinoma (HNSCC) | 6 | 5 |  |  | Treatment naïve |
| GSE234933 | Bill et al<sup>62</sup> | Head and neck squamous cell carcinoma (HNSCC) | 38 | 11 |  |  | not specified |
| GSE154778 | Lin et al<sup>63</sup> | Pancreatic ductal adenocarcinoma (PDAC) | 3 |  |  |  | not specified |
| GSE156405 | Lee et al<sup>64</sup> | Pancreatic ductal adenocarcinoma (PDAC) | 3 | 3 |  |  | Treatment naïve/ CTX/ IO/ Other |
| GSE197177 | Zhang et al<sup>65</sup> | Pancreatic ductal adenocarcinoma (PDAC) | 3 | 3 |  |  | Treatment naïve |
| GSE214295 | N/A | Pancreatic ductal adenocarcinoma (PDAC) | 3 |  |  |  | Treatment naïve |
| GSE231535 | Oh et al<sup>66</sup> | Pancreatic ductal adenocarcinoma (PDAC) | 2 |  |  |  | not specified |
| GSE183916 | Lenos et al<sup>67</sup> | Colorectal carcinoma (CRC) | 1 | 3 |  |  | not specified |
| GSE167297 | Jeong et al<sup>68</sup> | Gastric Adenocarcinoma (GAC) | 5 |  |  |  | not specified |
| GSE234129 | Wang et al<sup>69</sup> | Gastric Adenocarcinoma (GAC) | 3 | 2 |  |  | Treatment naïve |
| GSE223063 | Krishna et al<sup>70</sup> | Glioblastoma (GBM) | 3 |  |  |  | not specified |
| GSE224090 | N/A | Glioblastoma (GBM) | 2 |  |  |  | not specified |
| GSE235676 | Mei et al<sup>71</sup> | Glioblastoma (GBM) | 18 |  |  |  | Treatment naïve/ IO/ TT |

### Three distinct DC subpopulations exist in human cancer

The curated Myeloid APC atlas contains a total of 498,023 cells, including 29,887 DCs from 589 samples (**Figure 1a-e**). Most myeloid APC were tissue-resident macrophages (*FCGR1A*^+^*, CD68*^+^*, CSF1R*^+^*, TREM2*^+^*, SPP1*^+^*, C1QA, C1QB, and C1QC*), with smaller populations of classical (*CD14*^+^) and non-classical (*FCGR3A*^+^) monocytes (*LYZ*^+^*, S100A8*^+^*, S100A9*^+^) and DC (**Figure 1b** and **Supplementary Figure 1**). Distinct cDC1 (*CLEC9A*^+^*, XCR1*^+^*, BATF3*^+^*, IRF8*^+^), cDC2 (*CD1c*^+^*, FCER1A*^+^*, CLEC10A*^+^) and mregDC (*LAMP3*^+^*, CD274*^+^*, CCR7*^+^) clusters could be identified in this combined dataset. DC3 gene signatures^4,5^ (**Supplementary Table 1**), which are reported to overlap with both cDC2 and monocytes^5–7,13^ were most highly enriched within the classical monocyte population (**Figure 1f, Supplementary Figure 2a**) and did not align with a discrete cluster. Similarly, signatures for MoDC (‘DC4_FCGR3A’^4^) and reported pro-inflammatory cDC2/DC3-like populations^3^ (‘DC2_IL1B’, ‘DC2_FCN1’) were also most highly enriched within the macrophage and monocyte compartments (**Supplementary Figure 2b-d**).

We next extracted and re-clustered the DC populations separately and confirmed the identity of the three major DC populations conserved across tumour sites and datasets as being cDC1,^5^ cDC2,^5^ and mregDC^14^ (**Figure 2a-c, Supplementary Table 1, 2**). While the DC3 signature^5^ showed some alignment with the cDC2 cluster we could not segregate clear cDC2 / DC3 subpopulations based on this signature (**Figure 2b**).

**Figure 2:**
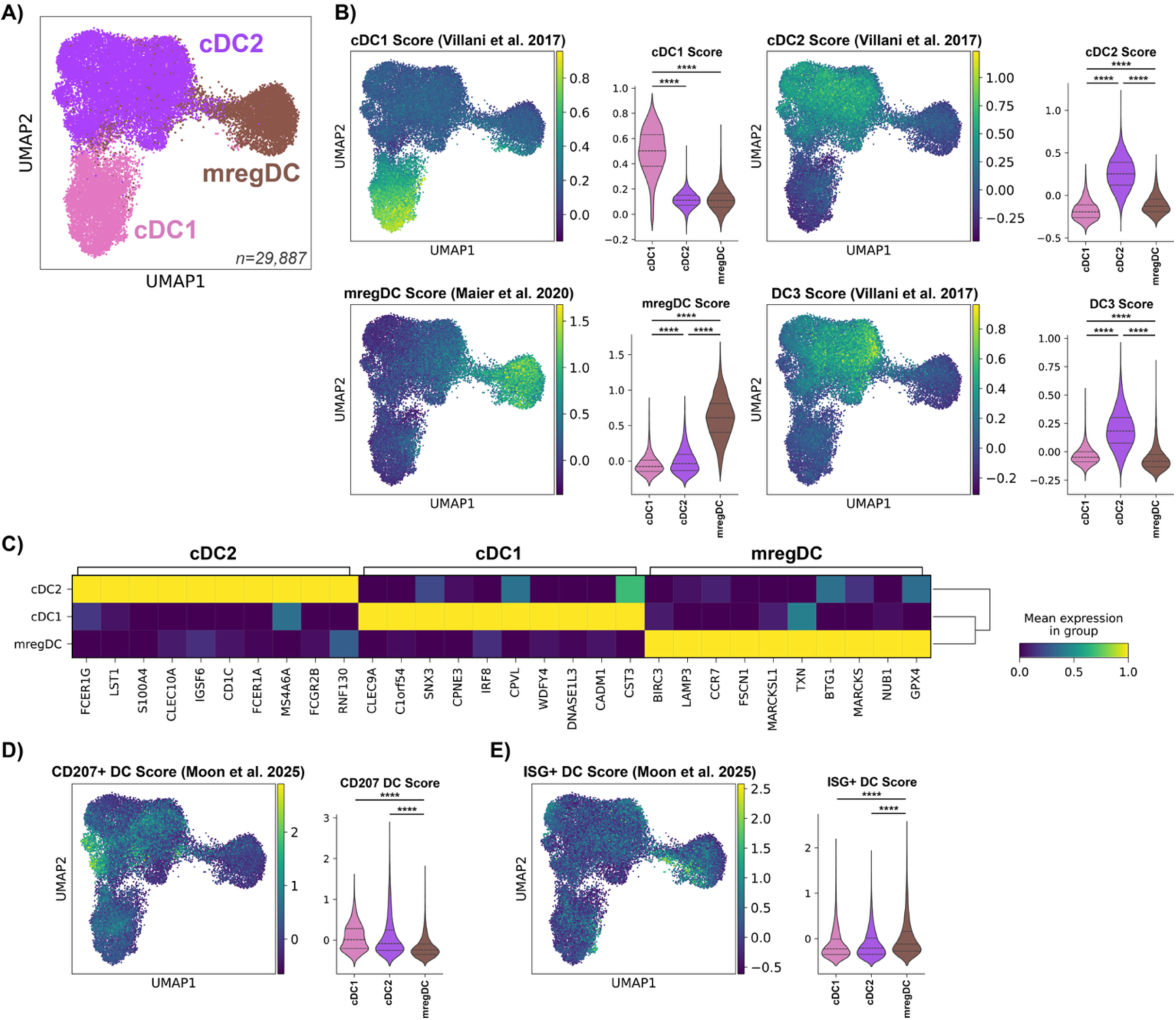
Alignment of DC signatures. **(A)** Analysis repeated on DC populations alone with BBKNN batch correction, and previous annotations confirmed with published signatures **(B)** from Maier et al^14^ and Villani et al^5^. **(C)** Differentially expressed genes (DEGs) between DC clusters were identified, portrayed in heatmap. DC signatures from Moon et al^1^ for CD207^+^ DC **(D)** and ISG^+^ DC **(E)** were overlaid on the atlas, and mean score per cell presented with violin plot grouped by subset. Enrichment scores compared between subsets using one-way ANOVA with Tukey’s multiple comparisons test. ****p<0.0001. Detailed statistics in Supplementary Table 3.

We also looked for the presence of the CD207^+^ DC and ISG^+^ DC that have been considered as maturation states that could potentially be adopted by multiple DC subsets^1^ (**Supplementary Table 1**). When projected onto the atlas, the CD207^+^ DC signature showed enrichment within a small proportion of the cDC2 compartment (**Figure 2d**) whilst the ISG^+^ DC signature was dispersed throughout the DC compartment (**Figure 2e**). There was, however, an area of heightened ISG enrichment within the mregDC region closest to cDC2, suggesting that ISG^+^ DC may represent a transitional state of cDC2 prior to the late maturation state of mregDC. We also note that type I interferon is reported to induce pleiotropic functions within the tumour microenvironment,^15^ and therefore, upregulation of the ISG program across populations likely represent DCs responding to these stimuli.

### Proportion of DC subsets varies by cancer and tissue type

Next, we examined the distribution of the myeloid APC cell types across the various cancers and tissues. In most of the included cancer types, DCs represent a minority (average of less than 15%) of total myeloid APCs (**Figure 3a,b**). However, in both primary MEL and NPC, DC proportion is higher (mean proportions of 48% and 32% respectively), with comparatively smaller proportions of both macrophages and monocytes seen in these cancers (**Supplementary Figure 1**).

**Figure 3:**
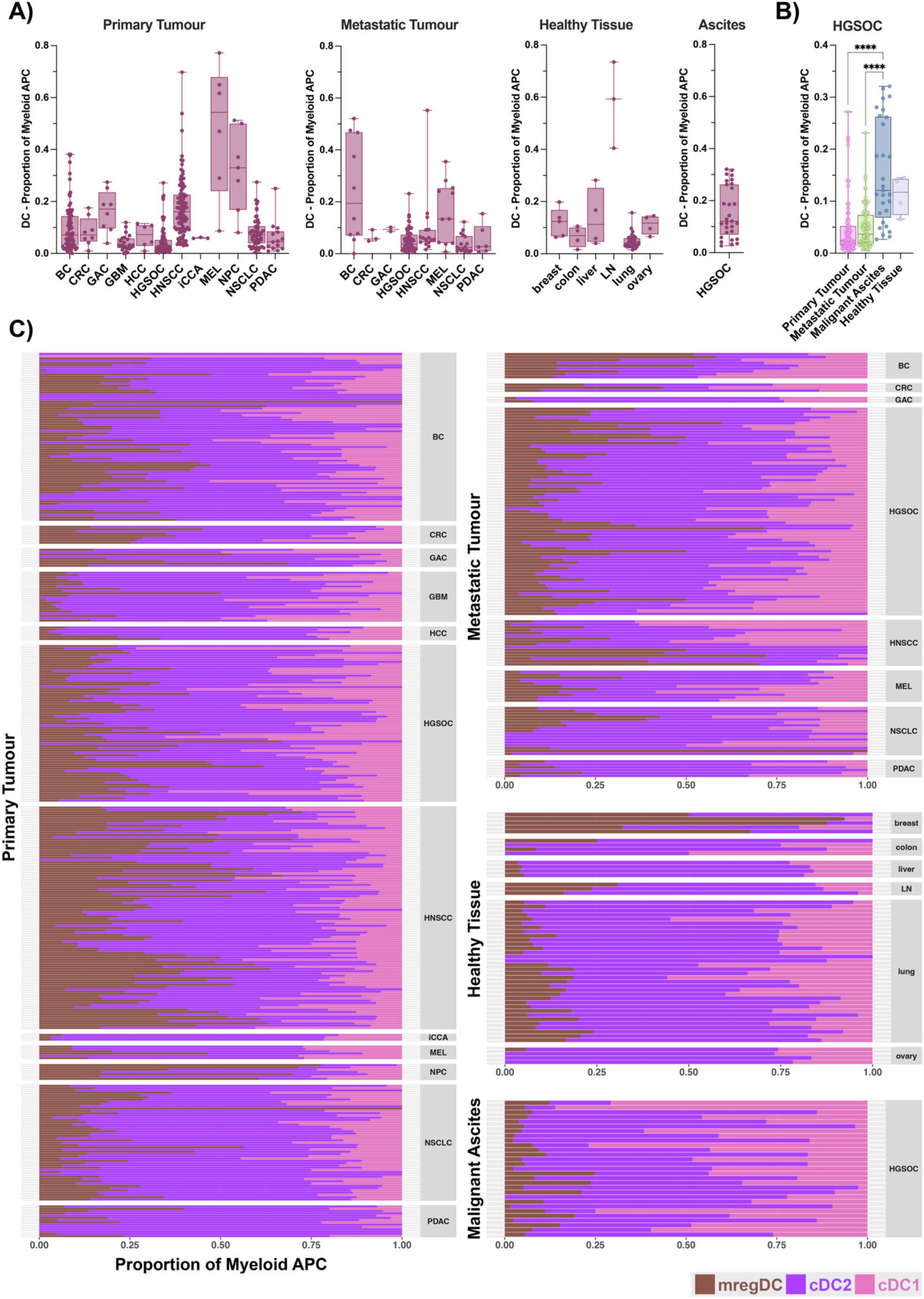
DC proportions in curated atlas. **(A)** Proportion of total DCs among all myeloid APCs is presented for each sample type. Box plot shows median with overlaid dots representing each individual sample. **(B)** Data from (A) shown for HGSOC samples only. Sample types compared using one-way ANOVA with Tukey’s multiple comparisons test. ****p<0.0001. Detailed statistics in Supplementary Table 3. **(C)** Proportion of each DC subset among total DCs is presented for primary, metastatic, and ascites samples along with healthy tissue as a stacked bar. Each bar represents a single sample, coloured by DC subset.

Aligning with previous studies,^3,16–18^ within primary tumours cDC2 represent the majority of DCs in most cancer types (**Figure 3c**). The exception was NPC, where in most samples, the majority of DCs were mregDCs, consistent with observations by Cheng et al^3^. Metastatic tumours showed similar proportions compared to their primary tumour counterparts, suggesting a conserved DC population at distant tumour sites. Interestingly, while only present for HGSOC, malignant ascites contained cDC1, cDC2 and mregDC (**Figure 3a-c**), with a significantly higher proportion of DCs (p<0.0001) compared to primary or metastatic HGSOC (**Figure 3b**). In most healthy tissues (excluding lymph node) DCs represented a minority of myeloid APCs, made up mostly of cDC2, with only low proportions of cDC1 and mregDC. The exception was healthy breast, where a high proportion of mregDC (average of over 60% of total DCs) was observed. To our knowledge this has not yet been reported, and no obvious reason was found within the literature to explain this, warranting further investigation to confirm the role of mregDCs in breast tissue.

### DC subset phenotype is highly conserved across tissues and cancers

Within the primary tumour samples of the myeloid APC atlas, each DC subset overlapped closely on the UMAP regardless of cancer type (**Figure 4a**). To further confirm conservation of DC subset phenotype across cancer types we calculated correlation between each sample-DC type pair on pseudo-bulked data (**Figure 4b,c**). Here, the same DC population tended to cluster together despite differences in cancer or tissue, with the majority of cDC1s falling in cluster 1 of the dendrogram, and the majority of cDC2s in cluster 2. While cluster 3 comprised mostly mregDCs, 59% of mregDCs were found within clusters 1 or 2, highlighting their close relationship to cDC1 and cDC2. This grouping by DC subset, as opposed to by tissue/cancer type, highlights conservation of DC subset phenotype across distant sites.

**Figure 4:**
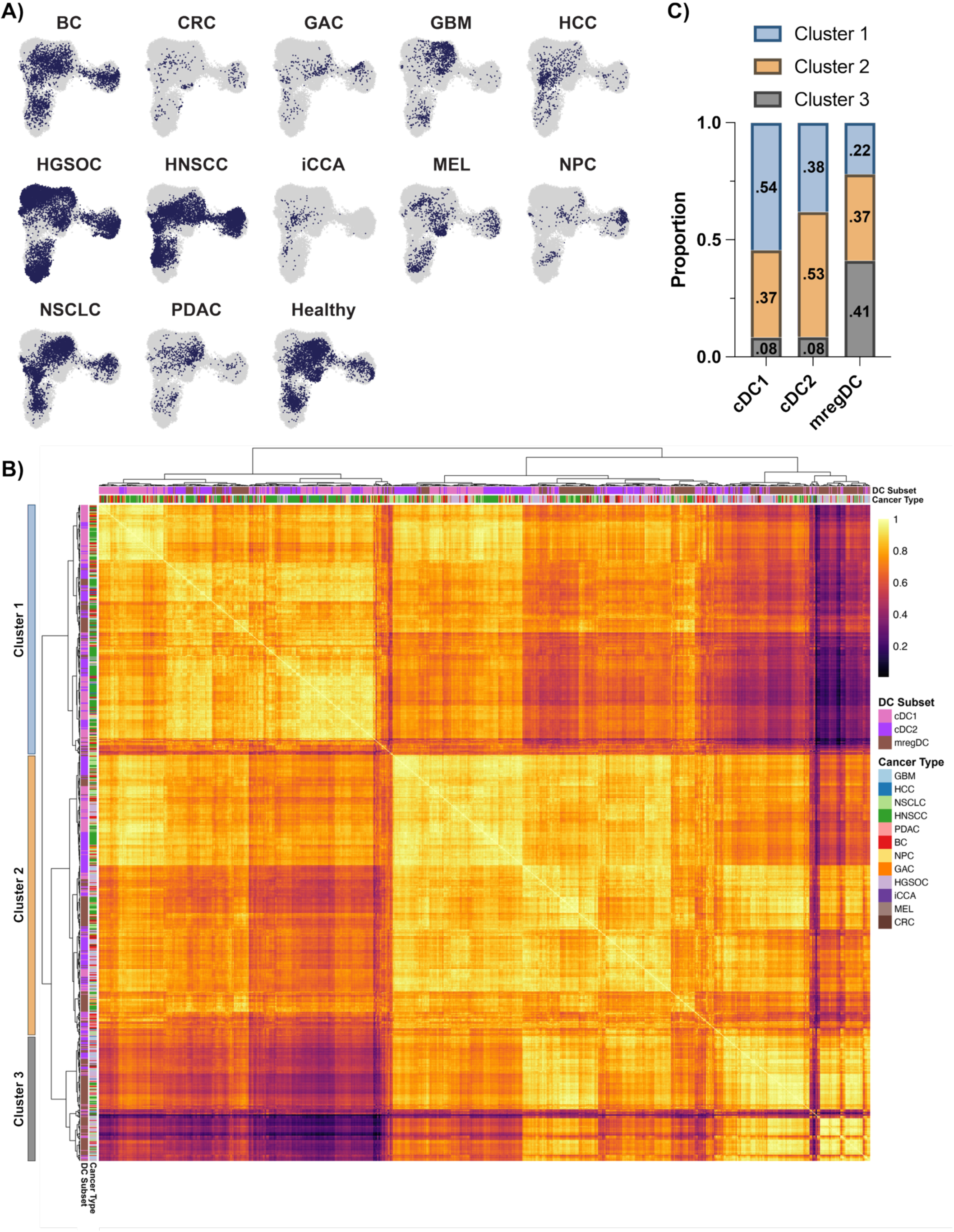
DC subsets are conserved across cancers and tissues. **(A)** UMAP was used to visualise curated DC atlas (including primary, metastatic and healthy tissues), split by cancer type. Healthy samples are all shown together as ‘Healthy’ **(B)** Pseudo bulking was performed on each DC subset from the curated myeloid APC atlas, here only primary tumour samples were considered. Pearson correlation was then performed between all sample-DC pairs. Heatmap shows correlation, cell subset and cancer type are indicated by legend. **(C)** Proportion of each DC subset among each of the three highest branches (clusters) in the predicted dendrogram from (B).

### Human tumour-associated cDC2 subsets align with cDC2A and cDC2B

Several studies have further segregated the cDC2 compartment within human tumours to as many as six different subtypes,^3,4^ but their alignment across datasets and whether they represent functionally distinct subsets or maturation states has not been established. To examine the degree of cDC2 compartmentalisation within the integrated myeloid APC atlas, we first extracted the cDC2 cluster and repeated the analysis and clustering pipeline at high resolution (**Figure 5a**, **Supplementary Figure 3a**). This identified 10 cDC2 subclusters varying in proportion across tissue samples and tumour types (**Figure 5b**). We next attempted to align the subclusters with previously annotated tumour associated cDC2 subsets (**Supplementary Table 1**).^3,4,19^ Signatures for ‘DC2_CXCR4’^3^ and ‘DC2_FCER1A’^4^ did not align with any particular cDC2 subcluster identified by unsupervised clustering analysis (**Figure 5c,d**). The ‘DC2_AREG’ signature^4^ was enriched across subclusters 1,3,5,7,8 whilst the ‘DC2_CXCL9’ signature^3^ was not enriched in any of the subclusters (**Supplementary Figure 3b,c**). Signatures for other previously described tumour-associated cDC2 subpopulations such as pro-inflammatory cDC2/DC3-like (‘DC2_IL1B’^3^ and ‘DC2_FCN1’^3^), DC3^4,5^, and MoDC (‘DC4_FCGR3A’)^4^ closely aligned with monocyte and macrophage populations (**Figure 1f**, **Supplementary Figure 2a-d**). Enrichment of the ‘DC2_FCN1’^3^ and ‘DC4_FCGR3A’^4^ signatures were also observed to a lesser extent within the cDC2 population but these were not aligned with any distinct cDC2 subcluster (**Supplementary Figure 3d-g**). However, the ‘DC2_IL1B’^3^ signature also aligned with cDC2 subcluster 1 (**Figure 5e**). Genes including *GPR183*, *NFKB1A*, *IL1B*, and *CD83* were amongst the top 20 DEGs for subcluster 1, suggesting this cDC2 subcluster is an inflammatory sub-state of cDC2 (**Supplementary Figure 3a**, **Supplementary Table 2**).^3^

**Figure 5:**
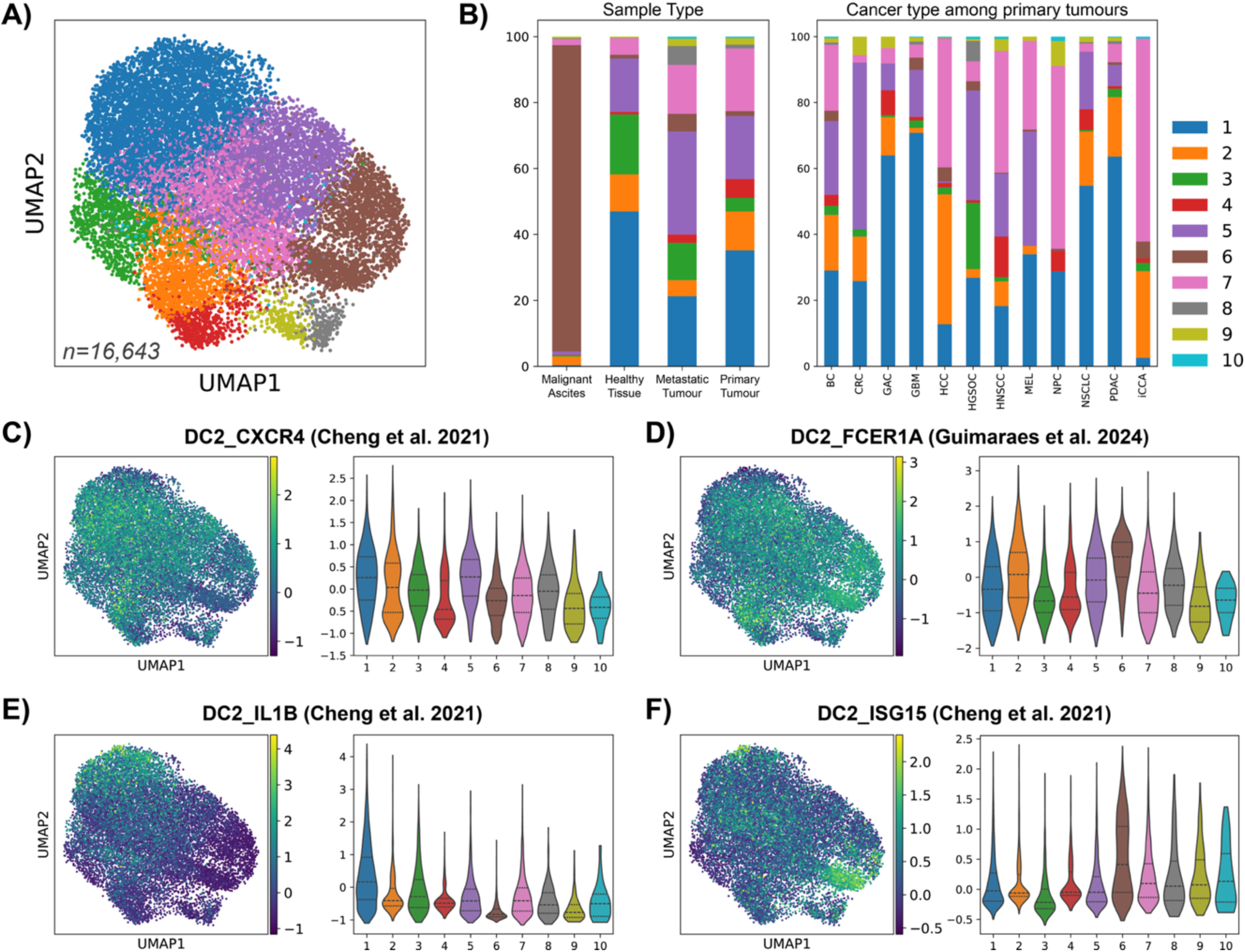
Subclustering cDC2 population. cDC2 cells were extracted from the myeloid APC atlas and analysis repeated. BBKNN batch correction was used for integration. **(A)** High resolution Leiden clustering identified 10 cDC2 clusters. **(B)** Proportion breakdown of cDC2 clusters is shown by sample type, and by cancer type (for primary tumours only)**. (C-F)** Enrichment for published signatures from Cheng et al^3^ and Guimaraes et al^4^ (as indicated) are shown overlaid on atlas and mean enrichment per cell plotted on violin plot grouped by cDC2 subcluster. Detailed statistics in Supplementary Table 3.

The cDC2 subcluster 6 was enriched for the ‘DC2_ISG15’ signature^3^ and was mostly comprised of samples from HGSOC malignant ascites (**Figure 5b,f**). Consistent with an interferon-induced activation state, cluster 6 was enriched in ‘interferon alpha/beta signalling’, ‘OAS antiviral response’, ‘Toll-like Receptor Cascades’ and ‘Interferon gamma signalling’ Reactome pathways (**Supplementary Table 2, Supplementary Figure 3h**). Subcluster 6 was also distinguished by a lack of *AREG* expression (**Supplementary Figure 3b**). This profile suggests that cDC2 in ascites may represent a more immunostimulatory subpopulation consistent with the ISG^+^ maturation state.

Notably, cDC2 subclusters 4, 8, 9, and 10 specifically aligned with signatures for ‘DC2_CD1A’3 and ‘DC2_CD207’4 in previous datasets (**Figure 6a,b**, **Supplementary Figure 4a**), unifying these cDC2 subtypes across different datasets as the CD207^+^ cDC2.

**Figure 6:**
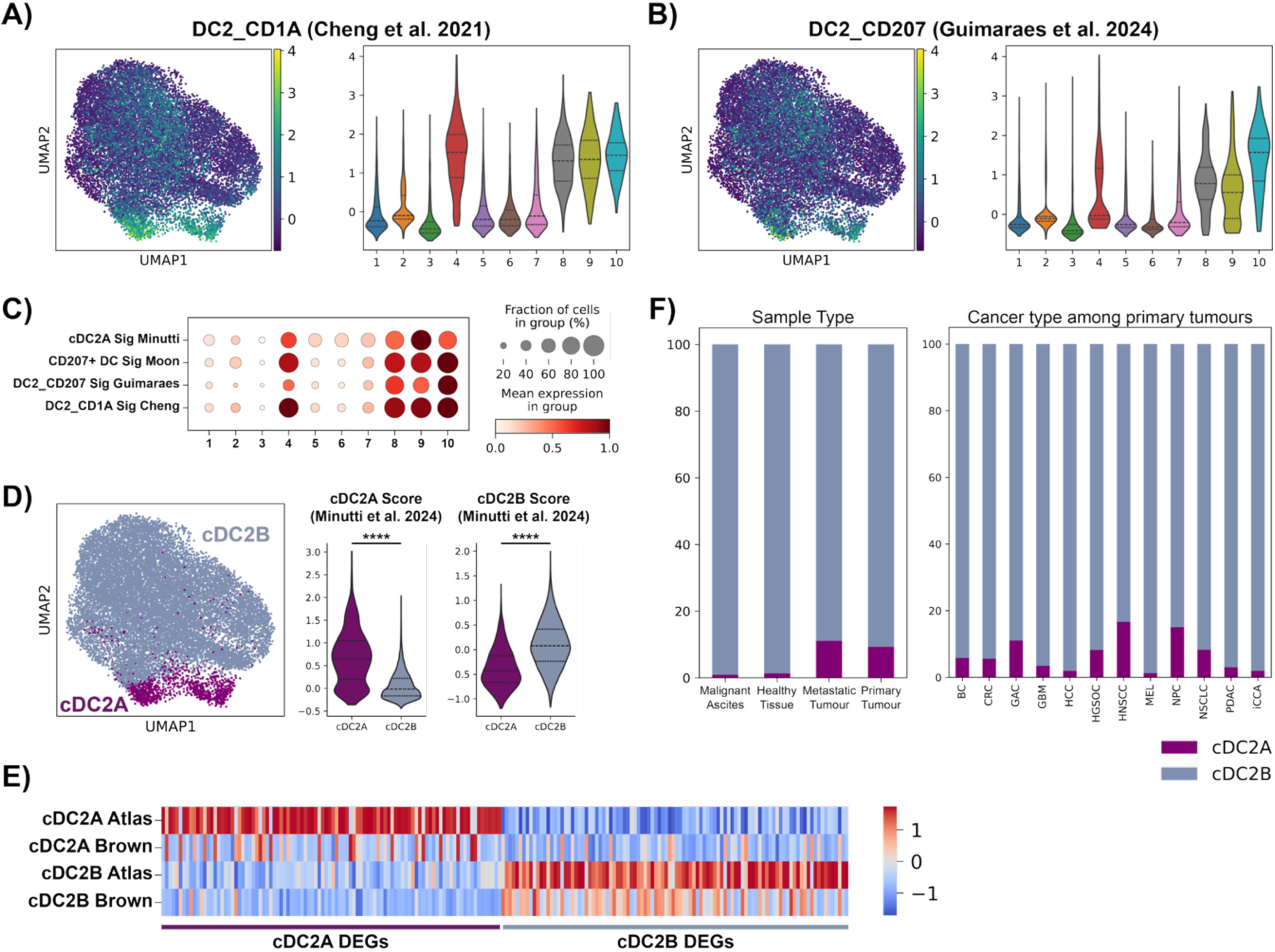
Tumour cDC2 can be divided into cDC2A and cDC2B. **(A,B)** Enrichment for published signatures from Cheng et al^3^ and Guimaraes et al^4^ (as indicated) are shown overlaid on atlas and mean enrichment per cell plotted on violin plot grouped by cDC2 subcluster. **(C)** Enrichment for published signatures^1,3,4,9^ shown as dot plot split by cDC2 subcluster. Dot size indicates percent of expressing cells, colour represents expression level. **(D)** cDC2 subclusters 4, 8, 9, 10 were identified as cDC2A and the remaining clusters cDC2B, these annotations are shown on UMAP and enrichment for published cDC2A/cDC2B signatures^9^ presented as violin plot. Enrichment scores compared between subsets using one-way ANOVA with Tukey’s multiple comparisons test, or via unpaired t-tests. ****p<0.0001. Detailed statistics for (A-D) in Supplementary Table 3. **(E)** cDC2A and cDC2B signatures were defined as the top 100 upregulated DEGs for each cell type. Average enrichment of these signature genes within cDC2A and cDC2B from human spleen data^20^ was compared to the cDC2A and cDC2B within the tumour atlas. Raw expression data was normalised prior to averaging per cell type; the four groups are then visually compared by plotting z-scaled averages for each gene via a heatmap. **(F)** Proportion breakdown of cDC2A/cDC2B shown by sample type, and by cancer type (for primary tumours only).

In murine and human spleen the cDC2 compartment can be separately subclassified into cDC2A and cDC2B subsets, that are proposed as ontogenetically distinct lineages with specialised functions.^19,20^ Some studies have incorrectly conflated the cDC2A/cDC2B division in lymphoid tissues with the cDC2/DC3 division in human blood^2,8^ (as human blood DC3 were initially designated as ‘CD1C_B’^5^). However, lymphoid tissue cDC2B are the more abundant cDC2 subset, align with circulating human CD1c^+^ DC (cDC2 and DC3),^20^ and are characterised by a proinflammatory phenotype.^20^ In contrast, cDC2A are absent from blood and express genes associated with immunomodulation and tissue repair.^20^ To understand how splenic cDC2A and B subsets relate to the cDC2 subclusters found in human cancer, tumour atlas cDC2 were scored with cDC2A and cDC2B signatures^19^ (**Supplementary Table 1**). The cDC2A signature closely aligned with subclusters 4, 8, 9, and 10, along with the CD207^+^ cDC2 signature, whilst the cDC2B signature aligned with the remaining clusters (**Figure 6c,d** and **Supplementary Figure 4b,c**).

Furthermore, human splenic cDC2A and cDC2B (as annotated by Brown et al^20^) aligned with signatures for the respective subsets in the tumour atlas (**Figure 6e**). Like their splenic counterparts, tumour associated cDC2B expressed genes typically associated with cDC2 including *CLEC10A* and *CLEC12A*, most toll-like receptors (TLRs), and were enriched in genes associated with inflammation (e.g., *IL1B, FCGR2A, GPR183, LGALS3* and *CXCL16*) and MHC class II processing (e.g., *CTSS* and *CD74*) (**Supplementary figure 4d,e, Supplementary table 2**). In contrast, cDC2A lacked expression of inflammatory TLRs and instead expressed *TLR10*, a human specific, anti-inflammatory TLR. cDC2A also expressed molecules involved in lipid antigen presentation (e.g., *CD1A* and *CD1E*) and regulation of mucosal immunity (e.g., *LTB, CCR6, IL22RA2* and *IL18*) (**Supplementary figure 4d**). Although cDC2A comprised the minor proportion of the cDC2 compartment, they were found in all primary tumour types and were enriched in both primary and metastatic tumour compared to healthy tissue and ascites (**Figure 6f**).

In summary, the atlas distilled cDC2 into two main subpopulations that were conserved across human cancer types: a minor cDC2A population enriched in tumour tissue defined by expression of *CD207* and *CD1A*, and a predominant cDC2B population defined by expression classical cDC2 associated genes such as *CLEC10A* and *CLEC12A*.

## Discussion

Single cell transcriptomics has revealed the complexity of the DC network with a multitude of subsets proposed as distinct ontogenetic lineages and/or maturation states. A deeper understanding of DC populations and their functions in the context of human tumours is critical to understanding how tumour immune responses are initiated or subverted, and how this can be exploited to develop more effective immunotherapies. Human pan-cancer myeloid atlases have been highly valuable in revealing the diversity of monocyte and macrophage populations, providing new biological insights and strategies for therapeutic targeting of these myeloid cells.^3,4^ They have also been useful in establishing the high degree of conservation among cDC1 and mregDC populations within human cancers and their broader alignment with equivalent subsets across tissue types and species. However, single-cell atlases have also highlighted the significant heterogeneity within the cDC2 compartment, with as many as six different cDC2 states reported with limited alignment between datasets.^3,4^ The rarity of DC within these datasets have been an additional barrier to resolving cDC2 heterogeneity. By careful curation and integration of suitable publicly available transcriptomic datasets, we have generated the most comprehensive pan- cancer single-cell atlas of DC in human tumours, comprised of 29,887 DCs (among a total of 498,023 myeloid APCs) from 589 samples. The atlas contains data from twelve solid tumour types, with both primary tumour samples, metastatic tumour samples (from eight cancer types), malignant ascites (from HGSOC), and associated healthy tissues (from six sites). This resource enables high resolution analysis for in depth characterisation of DCs across human cancer types.

Following unsupervised clustering, we applied the atlas to reconcile DC subsets with previously established signatures from human blood and various cancers. Consistent with earlier findings,^16^ cDC1, cDC2 and mregDC emerged as the three major DC populations, which were highly conserved among different human cancer types, albeit in different proportions. Further hierarchical clustering of the cDC2 compartment revealed 10 subclusters which did not fully align with the cDC2 subsets described in other datasets. Notably, we identified a small, distinct population of CD207^+^CD1A^+^ cDC2 corresponding to subsets previously annotated as ‘cDC2_CD1A’^3^ and ‘DC2_CD207’^4^, as well as to a condensed signature described in a recent review (CD207^+^ DC^1^). This finding consolidates CD207^+^CD1A^+^ DCs as a distinct cDC2 subtype across datasets. Our data demonstrate enrichment of CD207^+^CD1A^+^ DCs in tumours compared to healthy tissue, suggesting a potentially important role in the tumour microenvironment. A similar rare population of CD207^+^CD1A^+^ DC has also previously been described in several healthy human tissue types, with these cells being distinct from Langerhans cells (LC), which also express CD207 that encodes the C-type lectin receptor, Langerin.^21,22^

The origins and functions of tumour CD207^+^CD1A^+^ DC remain unclear, and a mouse equivalent tumour DC subset has not been identified due to species specific differences in *CD207* expression and the absence of *CD1A* in mice. However, we also showed that the core transcriptome signature of the CD207^+^CD1A^+^ DC population in human tumours aligns with another cDC2 subset found in human and mouse spleen, annotated as cDC2A.^19,20^ In both mouse and human, cDC2A are not present in circulation,^20^ and the common use of blood-derived signatures for annotation of DC populations in human tumour single-cell datasets may explain why this connection has not previously been made. The function of cDC2A is not yet fully understood, but they have been proposed to play a role in immune regulation.^20^ Genes involved in immune homeostasis in gut and lymphoid tissue, including *LTB*^23^ and *IL22RA2*^24^, were enriched in CD207^+^CD1A^+^ DCs (cDC2A) in human tumours, supporting a regulatory role. Enrichment of human specific *TLR10*, the only known TLR with anti-inflammatory functions,^25^ further supports a potential immunoregulatory role for tumour CD207^+^CD1A^+^ DCs, and expression of the human specific *CD1A* and *CD1E* suggests a role in the presentation of glycolipid antigens to T cells.^26^ Whether CD207^+^CD1A^+^ DC represent a unique DC lineage and/or a tissue induced maturation state of cDC2 remains to be established. *CD207* expression can be induced in monocytes and cDC2 by TGFβ *in vitro*, suggesting that CD207^+^CD1A^+^ DC could represent a maturation state of cDC2 and/or DC3^1^. However, their alignment with cDC2A suggests that tumour CD207^+^CD1A^+^ DCs may represent a separate ontogenetic lineage, similar to murine splenic cDC2A arising from a distinct bone marrow precursor.^19^ Overall, these findings highlight CD207^+^CD1A^+^ DCs as a distinct cDC2 subpopulation in human tumours that aligns with splenic cDC2A, and exhibits potential immunoregulatory functions, although their developmental origins and role in tumour immunity remain unresolved.

Aside from CD207^+^CD1A^+^ DCs, other subpopulations previously identified within the cDC2 cluster in human tumours include ‘cDC2_CXCL9’, ‘cDC2_IL1B’, ‘cDC2_FCN1’, ‘cDC2_ISG15’, and ‘cDC_CXCR4’^3^, along with ‘DC2_FCER1A’, ‘DC2_AREG’, ‘DC_CXCL8’, ‘DC3_CD14’ and ‘DC4_FCGR3A’^4^. However, when projected onto our atlas, these signatures were mostly found dispersed throughout the cDC2 population, rather than mapping to specific subclusters. Some of these populations, for example ‘cDC2_FCN1’^3^, ‘DC3_CD14’^4^, and ‘DC4_FCGR3A’^4^, were found enriched in the peripheral blood samples within these previous studies, and more closely resemble the blood DC3 subset^5^. Our atlas did not include peripheral blood samples, and overlaying the DC3 and related signatures did not identify a distinct DC3 population within tumour tissue. Rather, signatures for blood DC3 most closely aligned with tumour monocytes and macrophages and to a lesser extent with cDC2, highlighting the significant phenotypic overlap of DC3 with monocyte and cDC2 populations.

More broadly, we found the majority of the CD207⁻CD1A⁻ cDC2 (subclusters 1,2,3,5,6,7) most closely aligned with a signature classified as cDC2B in mouse and human spleen, which themselves align closely with human blood cDC2.^19^ When compared to tumour associated cDC2A (CD207^+^CD1A^+^) DC, tumour associated cDC2B were enriched in genes and pathways associated with TLR and inflammasome signalling, Fc receptors, and MHC class II processing, similar to their splenic and blood counterparts. Tumour cDC2B were also enriched in immunoregulatory pathways which may be reflective of the immunosuppressive TME. Within the cDC2B population we did find examples of maturation states, notably cDC2 subcluster 1, which was enriched in genes associated with inflammation and maturation, and aligned with the previously identified ‘cDC2_IL1B’ subpopulation.^3^ The cDC2 from malignant ascites samples clustered separately to tumour tissue within the cDC2B population as subcluster 6, and were distinguished by a more immunostimulatory phenotype characterised by enrichment of interferon signalling pathways. Aside from ascites samples, we found the greatest enrichment of the ISG signature within mregDC, providing evidence to support a relationship between these DC states in humans, as previously described in mice.^27,28^

In summary, our curated single-cell myeloid APC atlas confirms the major DC subsets present in human tumours and reveals an unrecognised alignment between CD207^+^CD1A^+^ cDC2 and cDC2A, with these cells exhibiting features suggestive of immunoregulatory roles. This atlas provides a valuable resource for future studies into myeloid APC diversity, phenotype, and function in human tumours.

### Study Limitations

Due to our focus on APCs, we included only HLA-DR expressing cells (myeloid APCs) in our atlas. As such, our atlas does not include the entire myeloid compartment, excluding non-HLA-DR expressing populations such as myeloid derived suppressor cells (MDSC) and mast cells.

## Methods

### Identification of datasets

We screened the DISCO database^12^ for relevant datasets and manually included additional datasets from select highly cited and recent studies not yet indexed on DISCO. Inclusion criteria were established to ensure consistent dataset selection, as summarised below:

*Dataset inclusion criteria:*

1. Published between 2019 and 2024 (inclusive).
2. scRNA-seq of tumour and/or related specimens from patients with diagnosed cancer (or associated healthy controls).
3. Solid tumours only.
4. Primary patient cells only (any cell lines, organoids, xenografts, or ex vivo treated samples excluded).
5. Generated using 10x technology (either 3’ or 5’ 10x sequencing).
6. Included all myeloid cells (samples pre-sorted for specific populations excluded).

Using these criteria, 57 datasets were initially identified for provisional inclusion in the atlas. Each dataset was then downloaded and further assessed for eligibility. Several datasets were excluded at this stage due to data issues (as discussed in Xu et al^29^), including those that are uploaded in inaccessible formats, contained incomplete data, lacked appropriate cell metadata or detailed methods regarding sample preparation, or had insubicient cell numbers or other signs of poor-quality sequencing. Following this round of exclusion, 36 datasets remained for inclusion in the atlas, covering a total of 12 major cancer types (summarised in **Table 1** and **Figure 1**). One dataset (GSE215120) included samples from both acral and cutaneous melanoma that were merged into one category, melanoma, for consistency. Similarly, as breast cancer subtypes were not specified in all datasets, all subtypes, including triple-negative breast cancer (TNBC), BRCA1^+^ TNBC, ER^+^ breast cancer, HER2^+^ breast cancer, ductal carcinoma in situ (DCIS), and invasive ductal carcinoma (IDC) were combined as ‘breast cancer’ (subtype retained in object metadata where available). Metastatic samples were included for HGSOC, MEL, BC, NSCLC, CRC, HNSCC, PDAC, and GAC, and these came from a variety of distant sites (details recorded in atlas metadata). Data from healthy tissue from breast, colon, liver, lung, lymph nodes, and ovary, taken from either healthy patients or adjacent non-tumour sites, were included. We excluded other sample types present in the original datasets (such as peripheral blood) from the atlas.

### Dataset download and myeloid APC extraction

From identified datasets, publicly accessible post-alignment scRNA-seq data was downloaded from corresponding online repositories. Gene expression matrices and associated files were then manually inspected to exclude datasets that provided only processed counts or inadequate sample metadata. Each of the remaining 36 datasets then underwent individual pre-processing, dimension reduction, and clustering to extract out the myeloid APCs using standard Seurat^30–34^ (v5.1.0) processing pipelines in R (v4.3.3), described below in detail (see **Supplementary Figure 5**).

The raw gene expression matrices were most often provided in the 10x Genomics format, consisting of separate barcodes, features/genes, and matrix files, or as 10X CellRanger hdf5 files. These were manually arranged into appropriate folder format and imported using the Read10X() or Read10X_h5() functions from the Seurat package in R, and Seurat objects generated for downstream analysis. In some cases, csv files, txt files, or Seurat objects were provided in lieu of 10X files, and these were read in directly. Provided sample metadata, including cancer type, sample type, patient id, dataset id, and a unique sample id/integration id were manually added to each sample. Note, any samples not biologically distinct (for example, two biopsies from the same tumour) were considered one sample for the purpose of batch correction and given the same integration id. All samples within a given dataset were then merged into one Seurat object.

Gene expression data was log-normalised using Seurat’s NormalizeData() function. The top 2,000 most variable genes across the dataset were identified for downstream dimensionality reduction and clustering using the FindVariableFeatures() function with the vst method. The data was then scaled using the ScaleData() function, and unwanted sources of variation, including the percentage of mitochondrial genes expressed and total RNA count per cell, were regressed out.

Principal component analysis (PCA) was performed using the RunPCA() function. Following this, nearest neighbours graph was generated and UMAP visualisation performed using the FindNeighbors() and RunUMAP() functions, with 40 dimensions. Each dataset was then clustered using default Louvain clustering with the FindClusters() function and 0.2 resolution. Canonical marker expression by each cluster was then visualised to determine which clusters contained myeloid APCs, and these preliminary APCs were extracted into a separate Seurat object. An example of this process is provided (**Supplementary Figure 6a,b**). Example canonical markers that were used to examine the data include, but not limited to: APC - *PTPRC* (CD45), *HLA-DR*; myeloid - *ITGAX* (CD11c), *ITGAM* (CD11b), *CD14*, and *FCGR3A* (CD16); cDC1 - *XCR1*, *CLEC9A*; cDC2/DC3 - *CD1C*, *FCER1A*; mregDC - *LAMP3*, *CCR7*; macrophage - *FCGR1A* (CD64), *CD68*, *CSF1R*; B cells - *CD19*; T cells - *CD3D*, *CD4*, *CD8A*.

The process described above was repeated for each of the 36 datasets individually (pipeline outlined in **Supplementary Figure 5**).

### Data processing and cleaning

All myeloid APCs were then concatenated into a single object. Only features present in all samples were retained (15,074 genes) to avoid bias during subsequent analyses. Only biologically distinct samples were considered unique, so in the case of multiple samples taken from the same tumour, these were considered the same sample for the remainder of the analysis. Any samples containing less than 100 myeloid APCs were excluded from further analysis.

Standard Scanpy pre-processing and analysis pipelines (python v3.12.3, Scanpy v1.10.2) were then employed to analyse the merged myeloid APC object (see **Supplementary Figure 5**). This was done in an iterative manner to continually remove contaminants until only myeloid APCs remained.

Firstly, gene expression data was normalised and log-transformed using scanpy.pp.normalize_total() and scanpy.pp.log1p() functions. The most highly variable genes were identified using the scanpy.pp.highly_variable_genes() function with default settings. Cell cycle scoring was performed using canonical markers for S and G2/M phases using the scanpy.tl.score_genes_cell_cycle() function and using cell cycling genes defined previously.^35,36^ These cell cycle scores (S phase and G2/M phase), along with effects of total RNA count per cell, and percentage of mitochondrial genes, were than regressed out using the scanpy.pp.regress_out() function. Data was scaled using the scanpy.pp.scale() function with values clipped at a standard deviation of 10. PCA was then performed using the scanpy.tl.pca() function with 60 dimensions. Harmony integration^37^ to correct for batch effects by sample was then performed using the scanpy.external.pp.harmony_integrate() function. Following integration, a nearest neighbours graph was generated using the scanpy.pp.neighbors() function with n_neighbors=10 and n_pcs=60. UMAP was then used to visualise the data with min_dist=0.3.

Leiden clustering was performed with the scanpy.tl.leiden() function and 0.2 resolution. Broad marker genes, like those used to identify myeloid APCs in each dataset, along with differentially expressed genes (DEG) between clusters (identified using the scanpy.tl.rank_genes_groups() function with Wilcoxon method), were then used to identify contaminating clusters within the object. Small clusters of contaminating T cells, NK cells, mast cells, keratinocytes, and epithelial cells were identified. These cells were then excluded from the earlier normalised and log- transformed object (prior to identification of highly variable genes). This analysis pipeline was then repeated until all contaminants removed and only confident myeloid DCs were retained within the DC clusters for further analysis. Following cleaning, the analysis pipeline and integration was then repeated on the final atlas. Clusters were annotated based on canonical markers, DEG genes, and published subset signatures as described in text. Note, here we employ Harmony integration for batch correction (**Supplementary Figure 7a-d)**, however comparison to mapping from scvi^38^ integration is shown (**Supplementary Figure 8a,b**), and annotations remain grouped together.

### DC and cDC2 Subclustering

The analysis pipeline was repeated on extracted DCs (cDC1, cDC2, and mregDC pooled) or cDC2s alone for visualisation and subclustering. For this, DC or cDC2 cells were extracted from the raw integrated atlas. Samples with less than 10 remaining cells or genes expressed in less than 10 cells were removed. Standard Scanpy analysis pipeline, as performed on the full atlas, was then repeated on the extracted cells from identification of highly variable genes to PCA. Batch balanced kNN (bbknn) batch correction^39^ using the scanpy.external.pp.bbknn() function was then used for integration, in place of Harmony and subsequent nearest neighbours, as it resulted in less distortion to the smaller dataset. Default Leiden clustering was then performed using the scanpy.tl.leiden() function and 0.8 resolution. For cDC2, subclustering of clusters was performed to reach adequate resolution. DEGs were identified using the scanpy.tl.rank_genes_groups() function with Wilcoxon method.

### Cell-cell correlation

DCs from primary tumours were extracted from the atlas and total gene expression per sample calculated. This was repeated separately for each of the three DC types. Pseudo-bulked data was normalised to 1e6. Correlation between all sample-DC pairs was then calculated using the cor() function (stats base package) with Pearson method in R. Correlations were plotted using the pheatmap package (v1.0.12).

### Pathways analysis

QIAGEN Ingenuity Pathway Analysis (IPA) software^40^ (QIAGEN Inc., https://digitalinsights.qiagen.com/IPA) was used to identify pathways from the Reactome database^41^ that were enriched within DEGs from clusters/cell types of interest. DEGs were identified as described previously, only those genes with adjusted p value < 0.05 and log fold change > 1.5 were included for IPA.

### Measuring enrichment of cDC2 signatures in human spleen

Signatures for cDC2A and cDC2B from the Myeloid APC atlas were defined as the top 100 upregulated DEGs from each cell type (genes were ordered by score and filtered for adjusted p value < 0.05 and log fold change > 0.58 (∼1.5-fold change)). Single-cell RNA sequencing data of human spleen^20^ was downloaded, and previous annotations of cDC2A (CLEC10⁻ cDC2) and cDC2B (CLEC10A^+^ cDC2) were retained. Raw expression data for the genes within the cDC2A and cDC2B signatures were extracted from the spleen^20^ and tumour (the Myeloid APC atlas) datasets. Two genes from the signatures that were not present within the spleen dataset were excluded.

Expression values were then normalised per gene, and average normalised expression calculated for each of the four groups (cDC2A and cDC2B from both spleen and tumour). Averages were then scaled (z-score) and presented as a heatmap.

### Visualisation and Statistics

All visualisation of the atlas and results were done using standard Scanpy (python) or Seurat (R) analysis and visualisation tools. Published signatures were overlaid on the atlas by generating an enrichment score for each cell using the scanpy.tl.score_genes() function. Enrichment scores were compared between groups using one-way ANOVA with Tukey’s multiple comparisons test (or unpaired t-tests for cases with only two groups) in GraphPad Prism 10. Detailed results from all statistical tests can be found in Supplementary Table 3. Stacked bar charts of number of samples/cells in the atlas, proportions within clusters following pseudo-bulking, and box plots of pooled proportions were generated using GraphPad Prism 10.

## Data and Code availability

The myeloid APC atlas will be accessible through an online interactive cellxgene viewer and as a downloadable .h5ad object upon publication. The code used in this manuscript will be released via a GitHub repository at the time of publication.

## Acknowledgements

KJR is supported by Mater Foundation and Tour de Cure. NR is supported by a University of Queensland Postgraduate Award Scholarship, Tour de Cure PhD support scholarship, and UQ Sam and Marion Frazer Top Up Scholarship in Immunotherapies. ZKT is supported by funding from the Children’s Hospital Foundation. XX is supported by the QLD Immunology Research Centre PhD Top-Up Scholarship.

## Author contributions

N.R. Z.K.T and K.J.R conceptualised and designed the study. N.R. and X.X. performed analysis.

N.R. wrote the manuscript with editing by Z.K.T and K.J.R. N.R. and N.T.Y. produced the interactive viewing platform. K.R. and Z.K.T. supervised the project.

## Supplementary Materials

**Supplementary Table 1: Published signatures.** Spreadsheet containing a summary of all published signatures overlaid on the myeloid APC atlas. Red text refers to genes not present in the atlas (not present in all datasets) and thus excluded from the signatures when overlaid.

**Supplementary Table 2: Differential gene expression analysis between DC subpopulations.** DEG analysis was performed using standard scanpy pipelines. DEG score, log fold change, p value, and adjusted p value is provided for all genes. Table include DEGs between cDC1, cDC2, and mregDC compared to all other DC, and then DEGs for all ten cDC2 clusters, cDC2A, and cDC2B groupings compared to all other cDC2.

**Supplementary Table 3: Detailed results from statistical analyses between groups.** Spreadsheet contains detailed results from all statistical analyses, including p values, between all contrasts for comparative statistics performed throughout manuscript. This includes one-way ANOVA and t-test comparisons done for Figure 1f, Figure 2b,d,e, Figure 3a, Figure 5c,d,e,f, Figure 6a,b,d, Supplementary Figure 2a,b,c,d, Supplementary Figure 3b,c,d,e,f,g, and Supplementary Figure 4a,b,c.

**Supplematary Figure 1:**
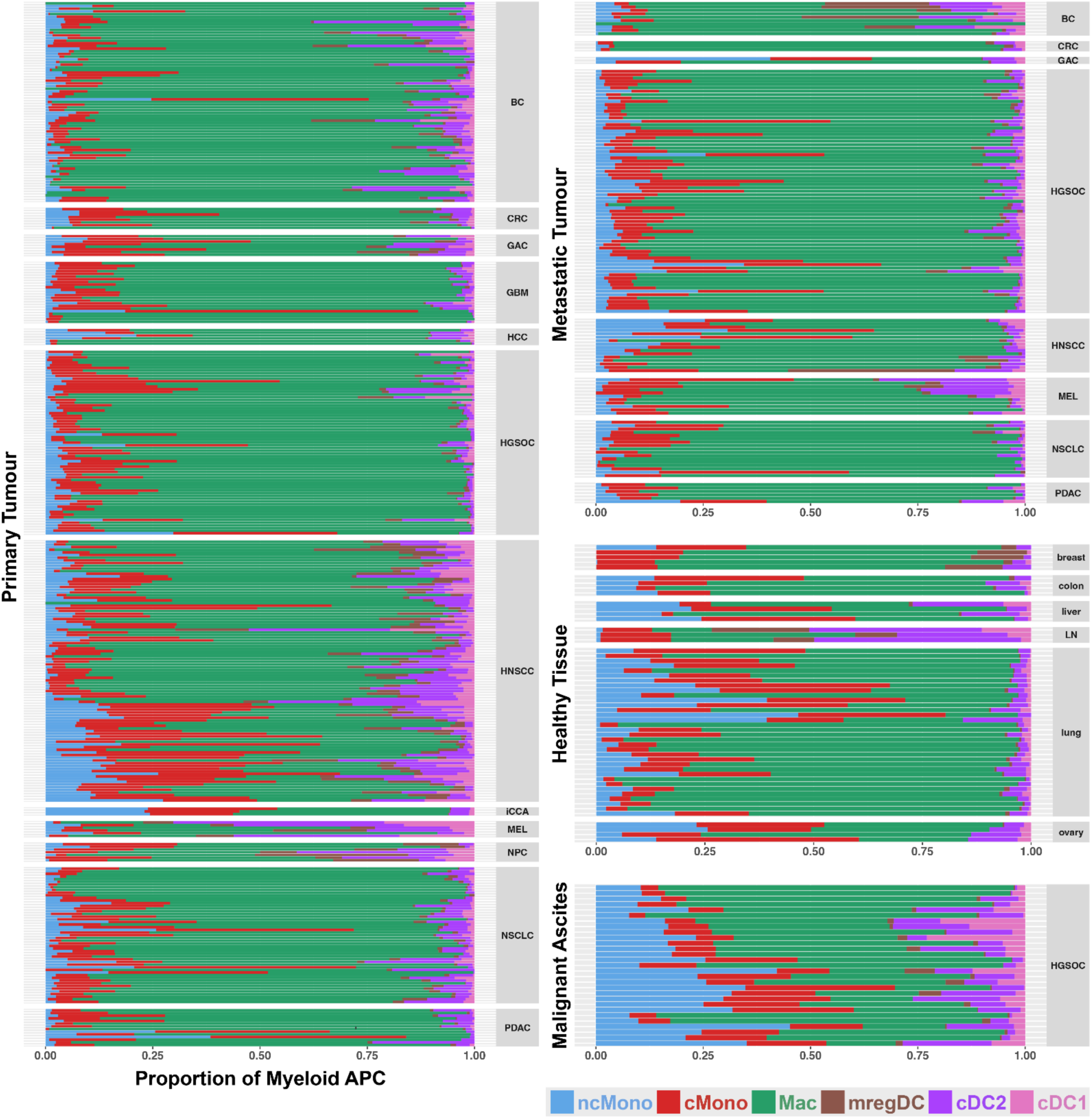
Cell type proportions in Myeloid APC atlas. Proportion of each cell type among total Myeloid APCs is presented for primary, metastatic, and ascites samples along with healthy tissue as a stacked bar. Each bar represents a single sample, coloured by cell type.

**Supplematary Figure 2:**
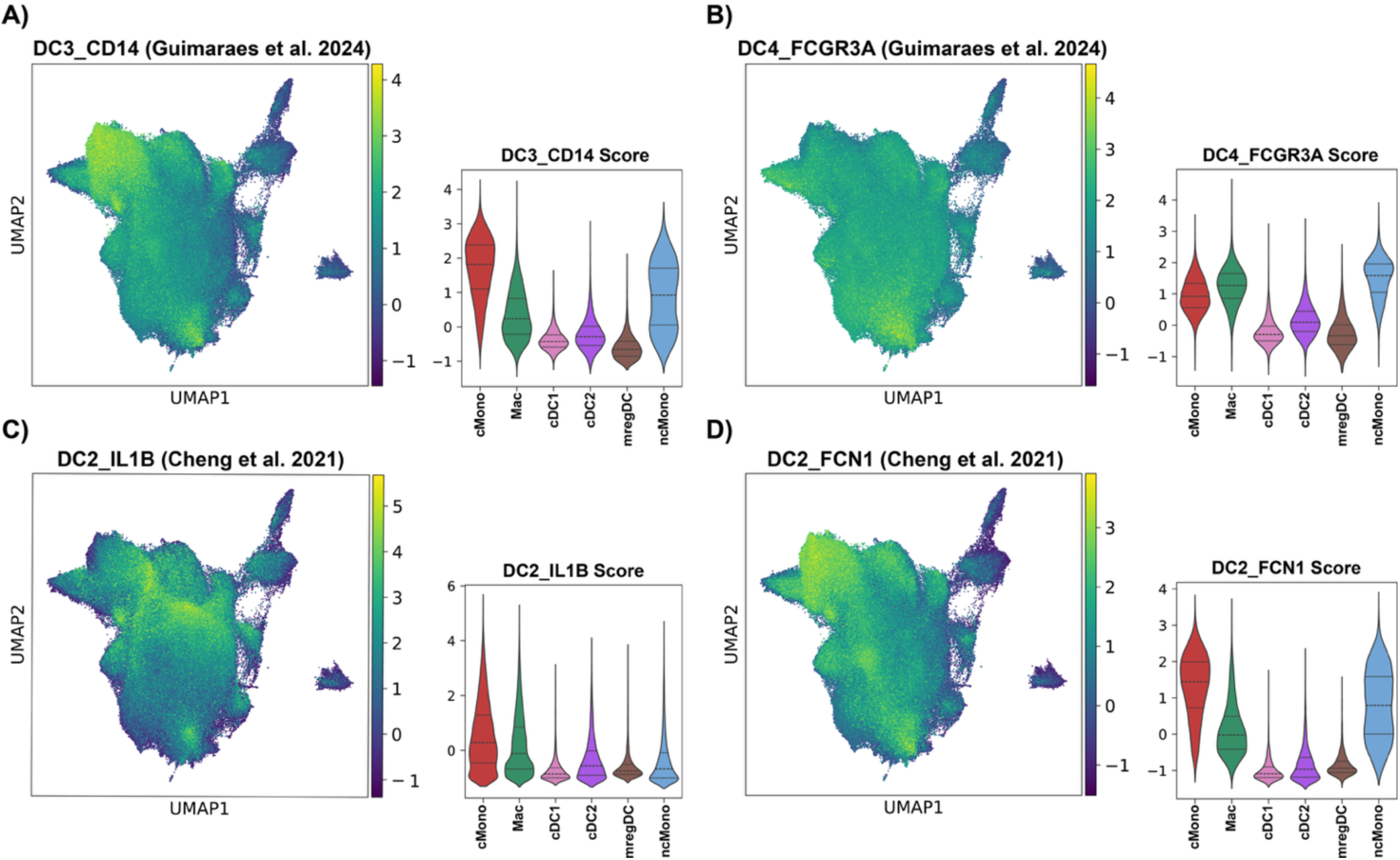
Published DC3, MoDC, and inflammatory DC signatures most highly enriched in monocyte and macrophage populations. Published signatures for **(A)** DC3 (‘DC3_CD14’)^4^, **(B)** MoDC (‘DC4_FCGR3A’)^4^, and **(C-D)** inflammatory cDC2/DC3-like cells (‘DC2_IL1B’ and ‘DC2_FCN1’)^3^ were overlaid on the myeloid APC atlas. Mean expression per cell is also shown with a violin plot, grouped by cell type. Detailed statistics in Supplementary Table 3.

**Supplementary Figure 3:**
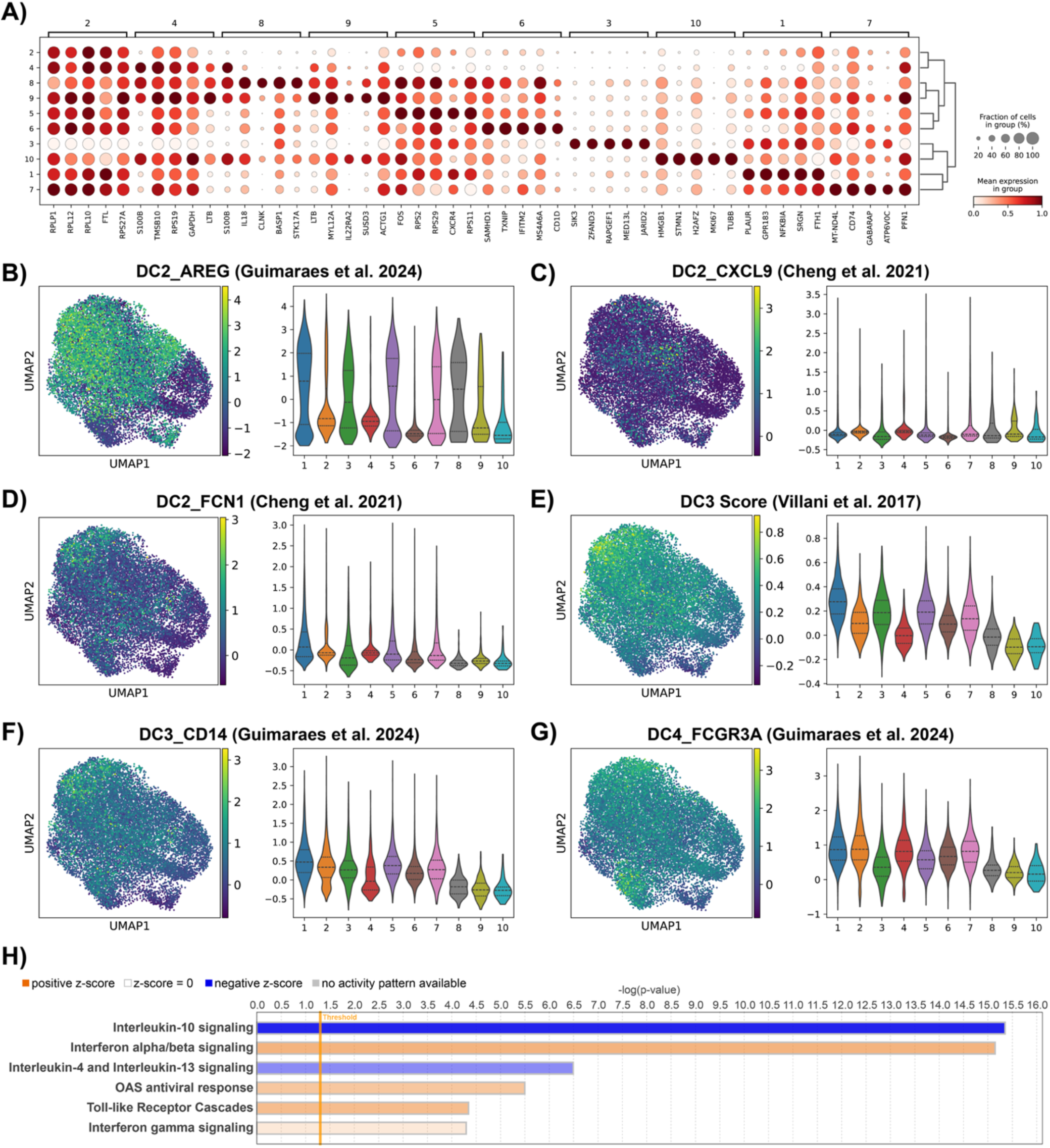
Subclustering cDC2 cells. **(A)** DEG genes were calculated for each cDC2 subcluster. Top 5 genes are shown with dot plot. Dot size indicates percent of expressing cells, colour represents expression level. **(B-G)** Enrichment for published signatures from Cheng et al^3^, Guimaraes et al^4^, and Villani et al^5^ (as indicated) are shown overlaid on atlas, and mean enrichment per cell plotted on violin plot grouped by cDC2 subcluster. Enrichment scores compared between subsets using one-way ANOVA with Tukey’s multiple comparisons test. Detailed statistics in Supplementary Table 3. **(H)** Pathway enrichment analysis was performed on DEGs using QIAGEN IPA software^40^, top Reactome pathways for cDC2 subcluster 6 shown. Colour indicates whether a pathway is predicted to be up- or down-regulated within the population, bar length indicates significance.

**Supplementary Figure 4:**
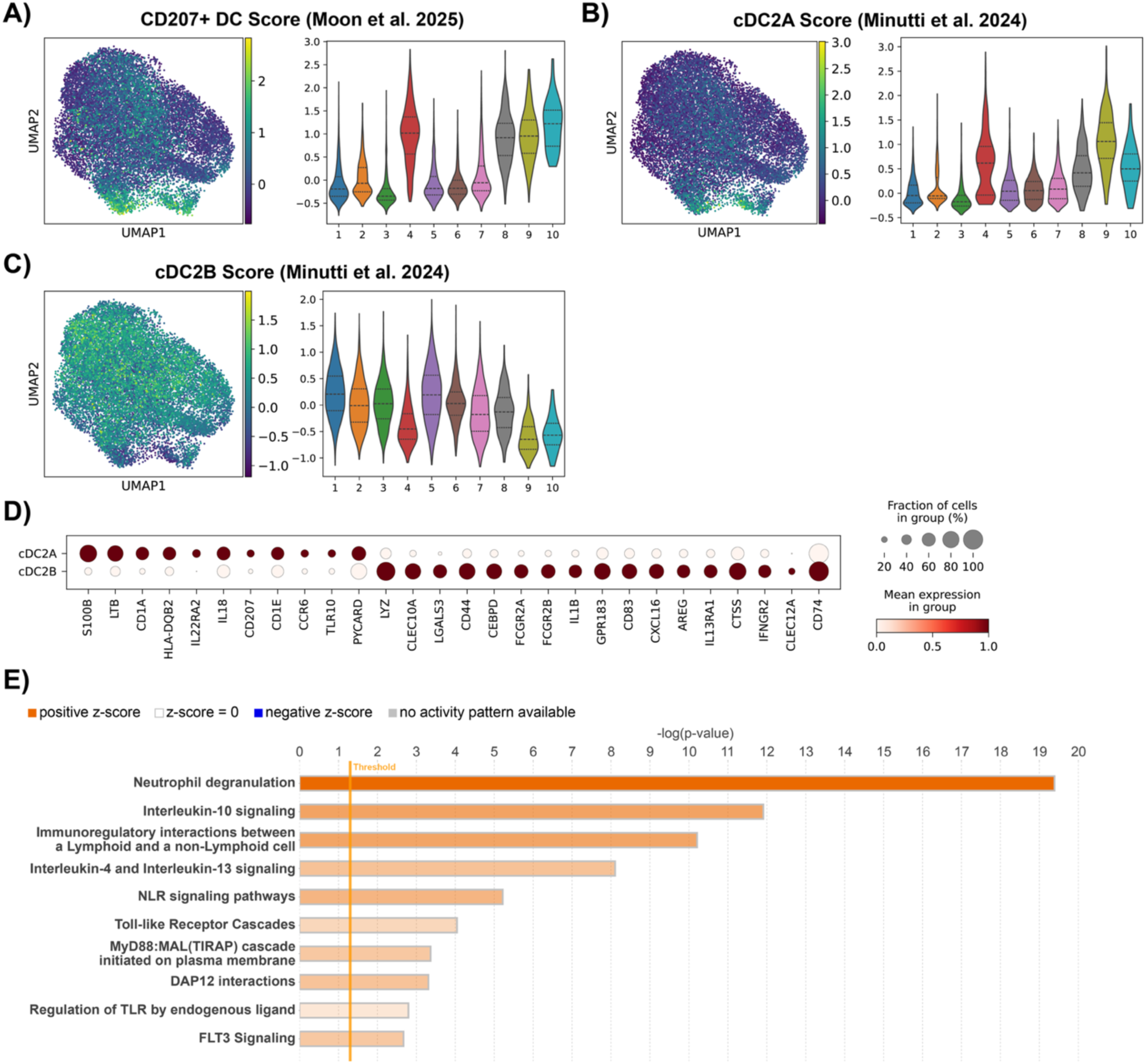
Tumour cDC2 can be divided into cDC2A and cDC2B. **(A-C)** Enrichment of cDC2 cells for CD207^+^ DC, cDC2A, and cDC2B signatures^1,19^ (as indicated) are shown overlaid on the UMAP and mean enrichment per cell plotted via violin plat grouped by cDC2 subcluster. Enrichment scores compared between subsets using one-way ANOVA with Tukey’s multiple comparisons test. Detailed statistics in Supplementary Table 3. **(D)** Select genes of interest are presented as a dot plot comparing cDC2A and cDC2B. Dot size indicates percent of expressing cells, colour represents expression level. **(E)** Pathway enrichment analysis was performed on DEGs for cDC2As and cDC2Bs using QIAGEN IPA software^40^, top Reactome pathways for cDC2B shown. Colour indicates whether a pathway is predicted to be up- or down-regulated within the population, bar length indicates significance.

**Supplementary Figure 5:**
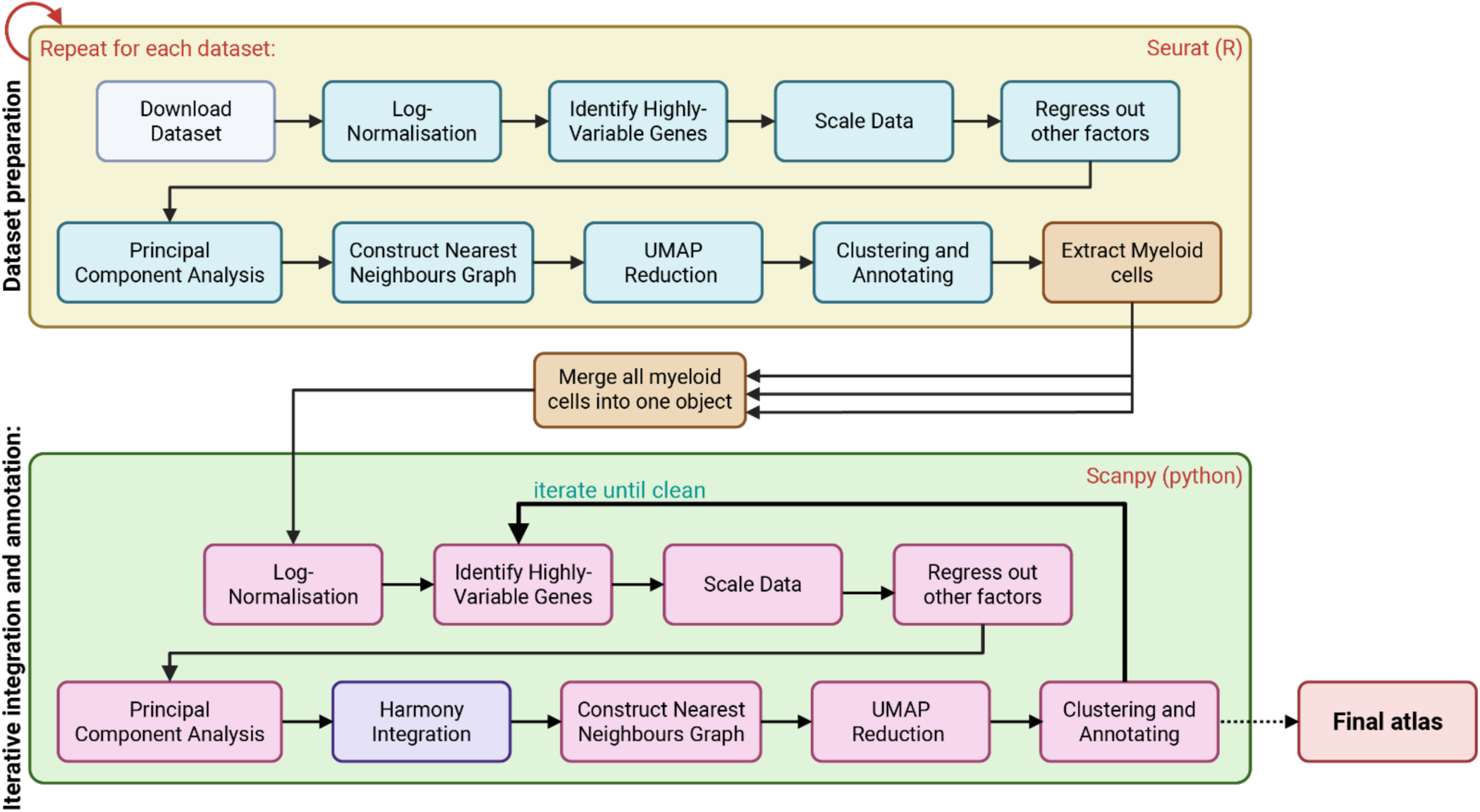
Preprocessing pipeline. Schematic shows the pipeline used to curate the single-cell myeloid APC atlas. Following download, each dataset was analysed using standard Seurat (R) pipeline. Myeloid APCs were then extracted from every dataset and merged into a single object. The merged object was analysed using standard Scanpy (python) pipeline. Harmony integration was used for batch correction. An iterative process was employed to ensure the final object contained only the cells of interest. Figure created with BioRender.com

**Supplementary Figure 6:**
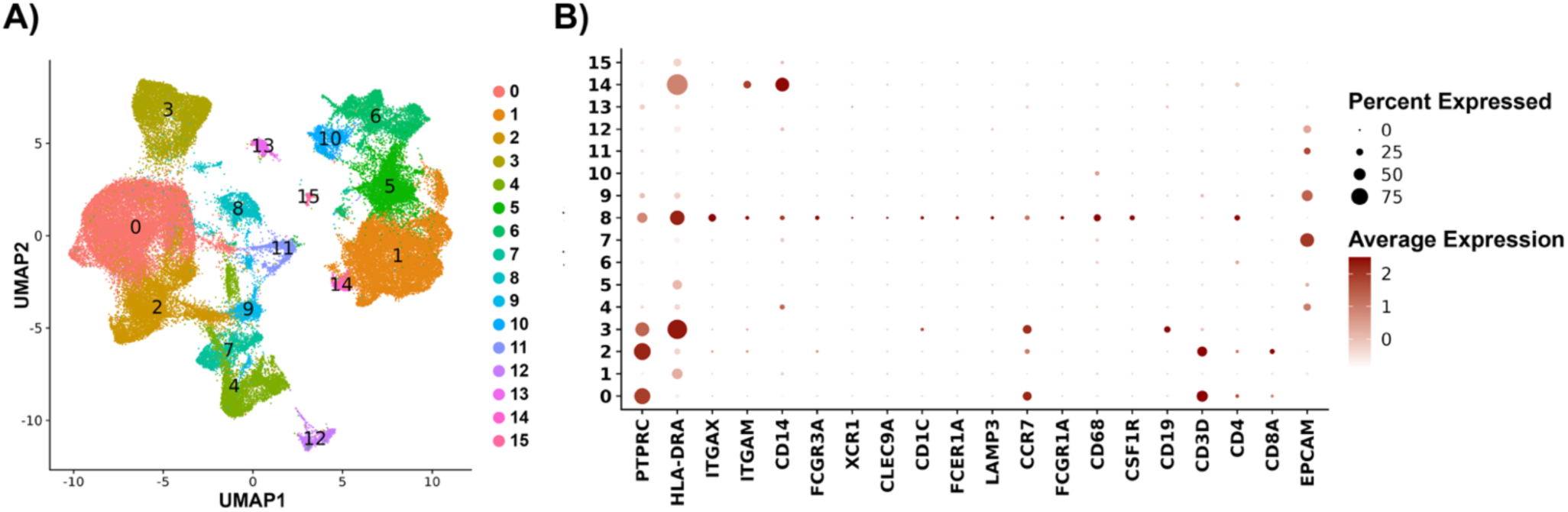
Extraction of myeloid APCs from public datasets. GSE225600 is presented as an example of the process repeated on all datasets included in the atlas. GSE225600 is a dataset containing eight samples, four patients diagnosed with breast cancer, with paired primary tumour and lymph node metastasis samples for each patient. Following download and QC, log-normalisation, variable gene identification, scaling, principal component analysis, and nearest neighbours graph generation, a UMAP dimension reduction and Louvain clustering was performed to visualise the data **(A)**. Canonical markers were then used to identify the clusters corresponding to myeloid APCs **(B)**. In this example cluster 8 was identified as myeloid APCs, this cluster was then extracted for downstream analyses. Dot size indicates percent of expressing cells, colour represents expression level.

**Supplementary Figure 7:**
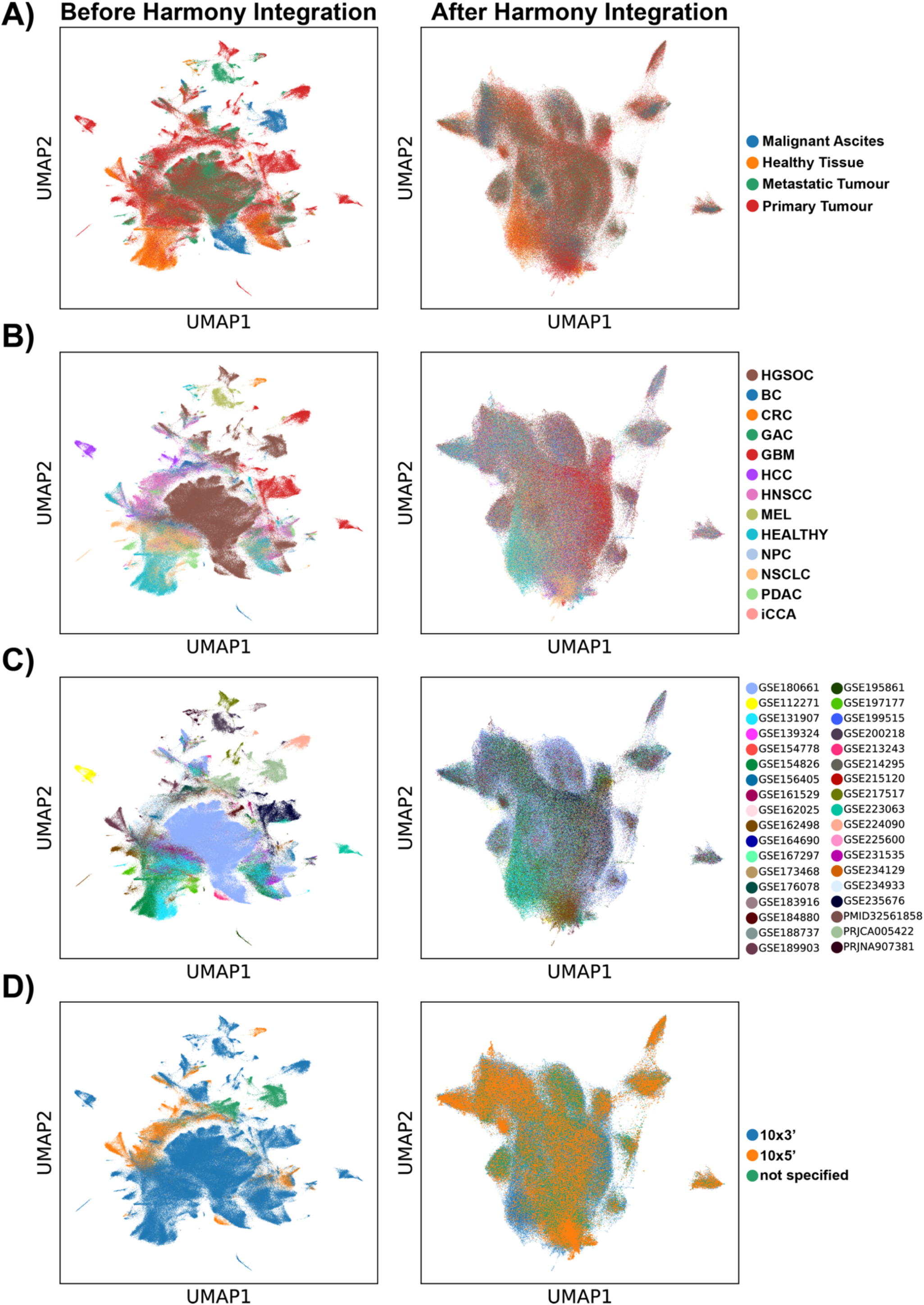
Harmony Integration on cleaned myeloid APC atlas. Harmony integration was used to integrate on a per sample basis. Plots show UMAP before and after Harmony integration, coloured by sample type **(A)**, cancer type **(B)**, original dataset **(C)**, and sequencing technology **(D)**.

**Supplementary figure 8:**
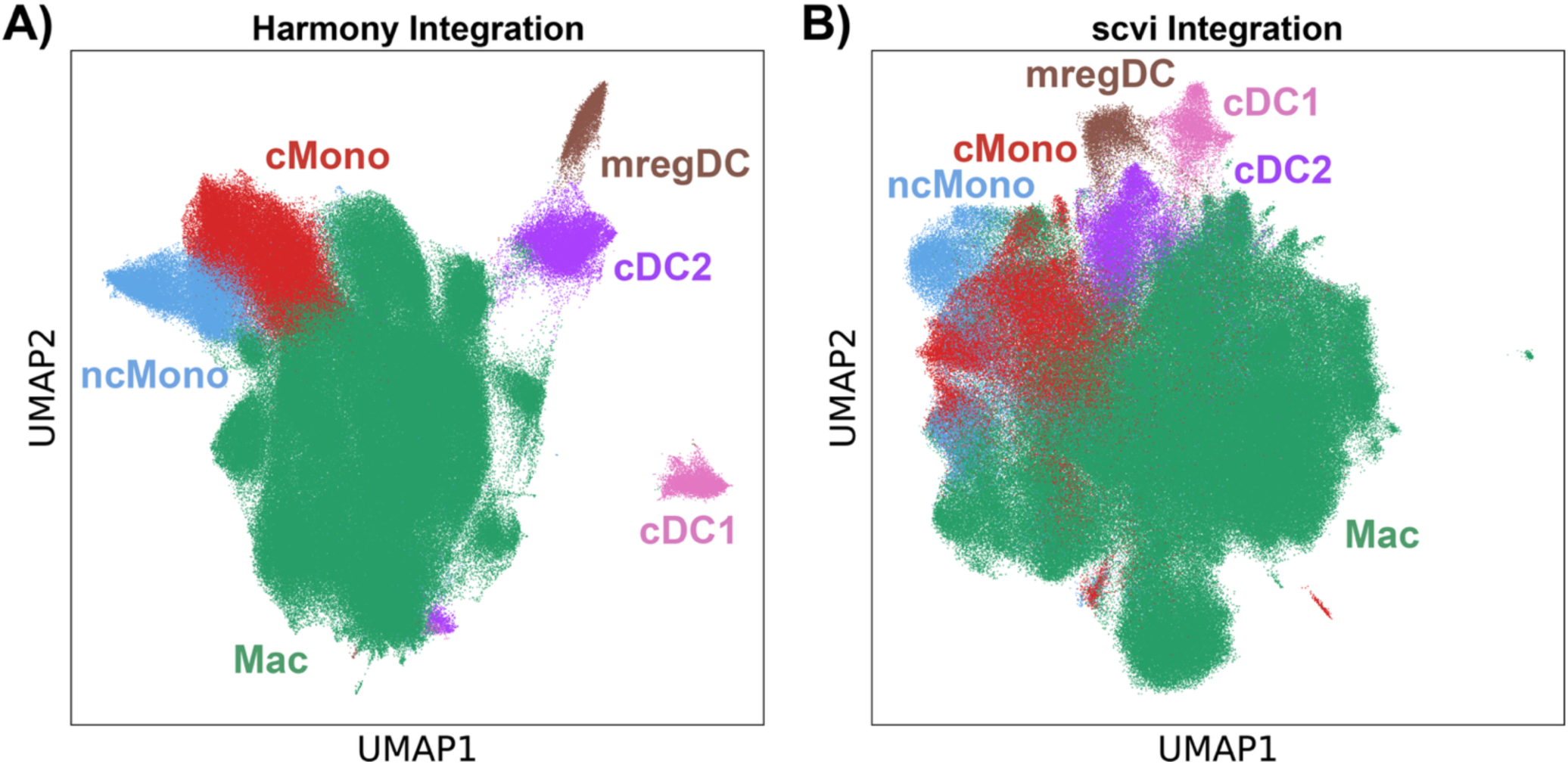
Comparison of integration methods. Integration was performed using both the Harmony^37^ **(A)** and scvi^38^ **(B)** method. Following subsequent cell type annotation on the Harmony object, annotations were transferred to the scvi object to confirm identified cell types formed distinct clusters irrespective of batch correction method.

## References

1. Moon CY, Belabed M, Park MD, Mattiuz R, Puleston D, Merad M. Dendritic cell maturation in cancer. Nat Rev Cancer. 2025;25(4):225–248. Published online 20250207. doi:10.1038/s41568-024-00787-3

2. Kvedaraite E, Ginhoux F. Human dendritic cells in cancer. Sci Immunol. 2022;7(70):eabm9409. Published online 20220401. doi:10.1126/sciimmunol.abm9409

3. Cheng S, Li Z, Gao R, et al. A pan-cancer single-cell transcriptional atlas of tumor infiltrating myeloid cells. Cell. 2021;184(3):792–809 e723. doi:10.1016/j.cell.2021.01.010

4. Guimaraes GR, Maklouf GR, Teixeira CE, et al. Single-cell resolution characterization of myeloid-derived cell states with implication in cancer outcome. Nat Commun. 2024;15(1):5694. Published online 20240707. doi:10.1038/s41467-024-49916-4

5. Villani AC, Satija R, Reynolds G, et al. Single-cell RNA-seq reveals new types of human blood dendritic cells, monocytes, and progenitors. Science. 2017;356(6335):eaah4573. doi:10.1126/science.aah4573

6. Bourdely P, Anselmi G, Vaivode K, et al. Transcriptional and Functional Analysis of CD1c(+) Human Dendritic Cells Identifies a CD163(+) Subset Priming CD8(+)CD103(+) T Cells. Immunity. 2020;53(2):335–352 e338. Published online 20200630. doi:10.1016/j.immuni.2020.06.002

7. Cytlak U, Resteu A, Pagan S, et al. Differential IRF8 Transcription Factor Requirement Defines Two Pathways of Dendritic Cell Development in Humans. Immunity. 2020;53(2):353–370 e358. Published online 20200730. doi:10.1016/j.immuni.2020.07.003

8. Liu Z, Wang H, Li Z, et al. Dendritic cell type 3 arises from Ly6C(+) monocyte-dendritic cell progenitors. Immunity. 2023;56(8):1761–1777 e1766. Published online 20230727. doi:10.1016/j.immuni.2023.07.001

9. Segura E, Touzot M, Bohineust A, et al. Human inflammatory dendritic cells induce Th17 cell differentiation. Immunity. 2013;38(2):336–348. Published online 20130124. doi:10.1016/j.immuni.2012.10.018

10. Bakdash G, Buschow SI, Gorris MA, et al. Expansion of a BDCA1+CD14+ Myeloid Cell Population in Melanoma Patients May Attenuate the Efficacy of Dendritic Cell Vaccines. Cancer Res. 2016;76(15):4332–4346. Published online 20160620. doi:10.1158/0008-5472.CAN-15-1695

11. Wculek SK, Cueto FJ, Mujal AM, Melero I, Krummel MF, Sancho D. Dendritic cells in cancer immunology and immunotherapy. Nat Rev Immunol. 2020;20(1):7–24. Published online 20190829. doi:10.1038/s41577-019-0210-z

12. Li M, Zhang X, Ang KS, et al. DISCO: a database of Deeply Integrated human Single-Cell Omics data. Nucleic Acids Res. 2022;50(D1):D596–D602. doi:10.1093/nar/gkab1020

13. Dutertre CA, Becht E, Irac SE, et al. Single-Cell Analysis of Human Mononuclear Phagocytes Reveals Subset-Defining Markers and Identifies Circulating Inflammatory Dendritic Cells. Immunity. 2019;51(3):573–589 e578. Published online 20190829. doi:10.1016/j.immuni.2019.08.008

14. Maier B, Leader AM, Chen ST, et al. A conserved dendritic-cell regulatory program limits antitumour immunity. Nature. 2020;580(7802):257-262. Published online 20200325. doi:10.1038/s41586-020-2134-y

15. Cheon H, Wang Y, Wightman SM, Jackson MW, Stark GR. How cancer cells make and respond to interferon-I. Trends Cancer. 2023;9(1):83–92. Published online 20221008. doi:10.1016/j.trecan.2022.09.003

16. Qian J, Olbrecht S, Boeckx B, et al. A pan-cancer blueprint of the heterogeneous tumor microenvironment revealed by single-cell profiling. Cell Res. 2020;30(9):745–762. Published online 20200619. doi:10.1038/s41422-020-0355-0

17. Vazquez-Garcia I, Uhlitz F, Ceglia N, et al. Ovarian cancer mutational processes drive site- specific immune evasion. Nature. 2022;612(7941):778-786. Published online 20221214. doi:10.1038/s41586-022-05496-1

18. Zheng X, Wang X, Cheng X, et al. Single-cell analyses implicate ascites in remodeling the ecosystems of primary and metastatic tumors in ovarian cancer. Nat Cancer. 2023;4(8):1138–1156. Published online 20230724. doi:10.1038/s43018-023-00599-8

19. Minutti CM, Piot C, Pereira da Costa M, et al. Distinct ontogenetic lineages dictate cDC2 heterogeneity. Nat Immunol. 2024;25(3):448–461. Published online 20240213. doi:10.1038/s41590-024-01745-9

20. Brown CC, Gudjonson H, Pritykin Y, et al. Transcriptional Basis of Mouse and Human Dendritic Cell Heterogeneity. Cell. 2019;179(4):846–863 e824. Published online 20191024. doi:10.1016/j.cell.2019.09.035

21. Bennett CL. Editorial: Faux amis: Langerin-expressing DC in humans and mice. J Leukoc Biol. 2015;97(4):621–623. doi:10.1189/jlb.5CE1014-481R

22. Bigley V, McGovern N, Milne P, et al. Langerin-expressing dendritic cells in human tissues are related to CD1c+ dendritic cells and distinct from Langerhans cells and CD141high XCR1+ dendritic cells. J Leukoc Biol. 2015;97(4):627–634. Published online 20141216. doi:10.1189/jlb.1HI0714-351R

23. Upadhyay V, Fu YX. Lymphotoxin signalling in immune homeostasis and the control of microorganisms. Nat Rev Immunol. 2013;13(4):270–279. doi:10.1038/nri3406

24. Martin JC, Beriou G, Heslan M, et al. Interleukin-22 binding protein (IL-22BP) is constitutively expressed by a subset of conventional dendritic cells and is strongly induced by retinoic acid. Mucosal Immunol. 2014;7(1):101–113. Published online 20130508. doi:10.1038/mi.2013.28

25. Hess NJ, Felicelli C, Grage J, Tapping RI. TLR10 suppresses the activation and differentiation of monocytes with effects on DC-mediated adaptive immune responses. J Leukoc Biol. 2017;101(5):1245–1252. Published online 20170224. doi:10.1189/jlb.3A1116-492R

26. Kim S, Cho S, Kim JH. CD1-mediated immune responses in mucosal tissues: molecular mechanisms underlying lipid antigen presentation system. Exp Mol Med. 2023;55(9):1858–1871. Published online 20230911. doi:10.1038/s12276-023-01053-6

27. Bosteels V, Marechal S, De Nolf C, et al. LXR signaling controls homeostatic dendritic cell maturation. Sci Immunol. 2023;8(83):eadd3955. Published online 20230512. doi:10.1126/sciimmunol.add3955

28. Ghislat G, Cheema AS, Baudoin E, et al. NF-κB-dependent IRF1 activation programs cDC1 dendritic cells to drive antitumor immunity. Sci Immunol. 2021;6(61). doi:10.1126/sciimmunol.abg3570

29. Xu X, Saxon J, Soon MSF, Lee CY, Tuong ZK. Data standards for single-cell RNA-sequencing of paediatric cancer. Clin Transl Immunology. 2025;14(5):e70033. Published online 20250523. doi:10.1002/cti2.70033

30. Butler A, Hoffman P, Smibert P, Papalexi E, Satija R. Integrating single-cell transcriptomic data across different conditions, technologies, and species. Nat Biotechnol. 2018;36(5):411–420. Published online 20180402. doi:10.1038/nbt.4096

31. Hao Y, Hao S, Andersen-Nissen E, et al. Integrated analysis of multimodal single-cell data. Cell. 2021;184(13):3573–3587 e3529. Published online 20210531. doi:10.1016/j.cell.2021.04.048

32. Hao Y, Stuart T, Kowalski MH, et al. Dictionary learning for integrative, multimodal and scalable single-cell analysis. Nat Biotechnol. 2024;42(2):293–304. Published online 20230525. doi:10.1038/s41587-023-01767-y

33. Satija R, Farrell JA, Gennert D, Schier AF, Regev A. Spatial reconstruction of single-cell gene expression data. Nat Biotechnol. 2015;33(5):495–502. Published online 20150413. doi:10.1038/nbt.3192

34. Stuart T, Butler A, Hoffman P, et al. Comprehensive Integration of Single-Cell Data. Cell. 2019;177(7):1888–1902 e1821. Published online 20190606. doi:10.1016/j.cell.2019.05.031

35. D C. Cell-Cycle Scoring and Regression. 2018. Accessed 4 June. https://notebook.community/theislab/scanpy_usage/180209_cell_cycle/cell_cycle

36. Tirosh I, Izar B, Prakadan SM, et al. Dissecting the multicellular ecosystem of metastatic melanoma by single-cell RNA-seq. Science. 2016;352(6282):189-196. doi:10.1126/science.aad0501

37. Korsunsky I, Millard N, Fan J, et al. Fast, sensitive and accurate integration of single-cell data with Harmony. Nat Methods. 2019;16(12):1289–1296. Published online 20191118. doi:10.1038/s41592-019-0619-0

38. Gayoso A, Lopez R, Xing G, et al. A Python library for probabilistic analysis of single-cell omics data. Nat Biotechnol. 2022;40(2):163–166. doi:10.1038/s41587-021-01206-w

39. Polański K, Young MD, Miao Z, Meyer KB, Teichmann SA, Park JE. BBKNN: fast batch alignment of single cell transcriptomes. Bioinformatics. 2020;36(3):964–965. doi:10.1093/bioinformatics/btz625

40. Krämer A, Green J, Pollard J, Jr., Tugendreich S. Causal analysis approaches in Ingenuity Pathway Analysis. Bioinformatics. 2014;30(4):523–530. Published online 20131213. doi:10.1093/bioinformatics/btt703

41. Milacic M, Beavers D, Conley P, et al. The Reactome Pathway Knowledgebase 2024. Nucleic Acids Res. 2024;52(D1):D672–D678. doi:10.1093/nar/gkad1025

42. Barrett T, Wilhite SE, Ledoux P, et al. NCBI GEO: archive for functional genomics data sets-- update. Nucleic Acids Res. 2013;41(Database issue):D991-995. Published online 20121127. doi:10.1093/nar/gks1193

43. Xu J, Fang Y, Chen K, et al. Single-Cell RNA Sequencing Reveals the Tissue Architecture in Human High-Grade Serous Ovarian Cancer. Clin Cancer Res. 2022;28(16):3590–3602. doi:10.1158/1078-0432.CCR-22-0296

44. Ren Y, Li R, Feng H, et al. Single-cell sequencing reveals effects of chemotherapy on the immune landscape and TCR/BCR clonal expansion in a relapsed ovarian cancer patient. Front Immunol. 2022;13:985187. Published online 20220928. doi:10.3389/fimmu.2022.985187

45. Hippen AA, Omran DK, Weber LM, et al. Performance of computational algorithms to deconvolve heterogeneous bulk ovarian tumor tissue depends on experimental factors. Genome Biol. 2023;24(1):239. Published online 20231020. doi:10.1186/s13059-023-03077-7

46. Biermann J, Melms JC, Amin AD, et al. Dissecting the treatment-naive ecosystem of human melanoma brain metastasis. Cell. 2022;185(14):2591–2608 e2530. doi:10.1016/j.cell.2022.06.007

47. Zhang C, Shen H, Yang T, et al. A single-cell analysis reveals tumor heterogeneity and immune environment of acral melanoma. Nat Commun. 2022;13(1):7250. Published online 20221125. doi:10.1038/s41467-022-34877-3

48. He M, Roussak K, Ma F, et al. CD5 expression by dendritic cells directs T cell immunity and sustains immunotherapy responses. Science. 2023;379(6633):eabg2752. Published online 20230217. doi:10.1126/science.abg2752

49. Pal B, Chen Y, Vaillant F, et al. A single-cell RNA expression atlas of normal, preneoplastic and tumorigenic states in the human breast. EMBO J. 2021;40(11):e107333. Published online 20210505. doi:10.15252/embj.2020107333

50. Wu SZ, Al-Eryani G, Roden DL, et al. A single-cell and spatially resolved atlas of human breast cancers. Nat Genet. 2021;53(9):1334–1347. Published online 20210906. doi:10.1038/s41588-021-00911-1

51. Tokura M, Nakayama J, Prieto-Vila M, et al. Single-Cell Transcriptome Profiling Reveals Intratumoral Heterogeneity and Molecular Features of Ductal Carcinoma In Situ. Cancer Res. 2022;82(18):3236–3248. doi:10.1158/0008-5472.CAN-22-0090

52. Liu YM, Ge JY, Chen YF, et al. Combined Single-Cell and Spatial Transcriptomics Reveal the Metabolic Evolvement of Breast Cancer during Early Dissemination. Adv Sci (Weinh). 2023;10(6):e2205395. Published online 20230103. doi:10.1002/advs.202205395

53. Gueguen P, Metoikidou C, Dupic T, et al. Contribution of resident and circulating precursors to tumor-infiltrating CD8(+) T cell populations in lung cancer. Sci Immunol. 2021;6(55). doi:10.1126/sciimmunol.abd5778

54. Kim N, Kim HK, Lee K, et al. Single-cell RNA sequencing demonstrates the molecular and cellular reprogramming of metastatic lung adenocarcinoma. Nat Commun. 2020;11(1):2285. Published online 20200508. doi:10.1038/s41467-020-16164-1

55. Leader AM, Grout JA, Maier BB, et al. Single-cell analysis of human non-small cell lung cancer lesions refines tumor classification and patient stratification. Cancer Cell. 2021;39(12):1594–1609 e1512. Published online 20211111. doi:10.1016/j.ccell.2021.10.009

56. Losic B, Craig AJ, Villacorta-Martin C, et al. Intratumoral heterogeneity and clonal evolution in liver cancer. Nat Commun. 2020;11(1):291. Published online 20200115. doi:10.1038/s41467-019-14050-z

57. Ma L, Heinrich S, Wang L, et al. Multiregional single-cell dissection of tumor and immune cells reveals stable lock-and-key features in liver cancer. Nat Commun. 2022;13(1):7533. Published online 20221207. doi:10.1038/s41467-022-35291-5

58. Liu Y, He S, Wang XL, et al. Tumour heterogeneity and intercellular networks of nasopharyngeal carcinoma at single cell resolution. Nat Commun. 2021;12(1):741. Published online 20210202. doi:10.1038/s41467-021-21043-4

59. Cillo AR, Kurten CHL, Tabib T, et al. Immune Landscape of Viral- and Carcinogen-Driven Head and Neck Cancer. Immunity. 2020;52(1):183–199 e189. Published online 20200107. doi:10.1016/j.immuni.2019.11.014

60. Kurten CHL, Kulkarni A, Cillo AR, et al. Investigating immune and non-immune cell interactions in head and neck tumors by single-cell RNA sequencing. Nat Commun. 2021;12(1):7338. Published online 20211217. doi:10.1038/s41467-021-27619-4

61. Quah HS, Cao EY, Suteja L, et al. Single cell analysis in head and neck cancer reveals potential immune evasion mechanisms during early metastasis. Nat Commun. 2023;14(1):1680. Published online 20230327. doi:10.1038/s41467-023-37379-y

62. Bill R, Wirapati P, Messemaker M, et al. CXCL9:SPP1 macrophage polarity identifies a network of cellular programs that control human cancers. Science. 2023;381(6657):515-524. Published online 20230803. doi:10.1126/science.ade2292

63. Lin W, Noel P, Borazanci EH, et al. Single-cell transcriptome analysis of tumor and stromal compartments of pancreatic ductal adenocarcinoma primary tumors and metastatic lesions. Genome Med. 2020;12(1):80. Published online 20200929. doi:10.1186/s13073-020-00776-9

64. Lee JJ, Bernard V, Semaan A, et al. Elucidation of Tumor-Stromal Heterogeneity and the Ligand-Receptor Interactome by Single-Cell Transcriptomics in Real-world Pancreatic Cancer Biopsies. Clin Cancer Res. 2021;27(21):5912–5921. Published online 20210823. doi:10.1158/1078-0432.CCR-20-3925

65. Zhang S, Fang W, Zhou S, et al. Single cell transcriptomic analyses implicate an immunosuppressive tumor microenvironment in pancreatic cancer liver metastasis. Nat Commun. 2023;14(1):5123. Published online 20230823. doi:10.1038/s41467-023-40727-7

66. Oh K, Yoo YJ, Torre-Healy LA, et al. Coordinated single-cell tumor microenvironment dynamics reinforce pancreatic cancer subtype. Nat Commun. 2023;14(1):5226. Published online 20230826. doi:10.1038/s41467-023-40895-6

67. Lenos KJ, Bach S, Ferreira Moreno L, et al. Molecular characterization of colorectal cancer related peritoneal metastatic disease. Nat Commun. 2022;13(1):4443. Published online 20220804. doi:10.1038/s41467-022-32198-z

68. Jeong HY, Ham IH, Lee SH, et al. Spatially Distinct Reprogramming of the Tumor Microenvironment Based On Tumor Invasion in Diffuse-Type Gastric Cancers. Clin Cancer Res. 2021;27(23):6529–6542. Published online 20210812. doi:10.1158/1078-0432.CCR-21-0792

69. Wang R, Song S, Qin J, et al. Evolution of immune and stromal cell states and ecotypes during gastric adenocarcinoma progression. Cancer Cell. 2023;41(8):1407–1426 e1409. Published online 20230706. doi:10.1016/j.ccell.2023.06.005

70. Krishna S, Choudhury A, Keough MB, et al. Glioblastoma remodelling of human neural circuits decreases survival. Nature. 2023;617(7961):599-607. Published online 20230503. doi:10.1038/s41586-023-06036-1

71. Mei Y, Wang X, Zhang J, et al. Siglec-9 acts as an immune-checkpoint molecule on macrophages in glioblastoma, restricting T-cell priming and immunotherapy response. Nat Cancer. 2023;4(9):1273–1291. Published online 20230717. doi:10.1038/s43018-023-00598-9

